# Functional integration of “undead” neurons in the olfactory system

**DOI:** 10.1101/623488

**Authors:** Lucia L. Prieto-Godino, Ana F. Silbering, Mohammed A. Khallaf, Steeve Cruchet, Karolina Bojkowska, Sylvain Pradervand, Bill S. Hansson, Markus Knaden, Richard Benton

## Abstract

Programmed cell death (PCD) is widespread during neurodevelopment, typically eliminating the surpluses of neuronal production. Employing the *Drosophila* olfactory system, we examined the potential of cells fated to die to contribute to circuit evolution. Inhibition of PCD is sufficient to generate many new cells that express neural markers and exhibit odor-evoked activity. These “undead” neurons express a subset of olfactory receptors that, intriguingly, is enriched for recent receptor duplicates and include some normally found in other chemosensory organs and life-stages. Moreover, undead neuron axons integrate into the olfactory circuitry in the brain, forming novel receptor/glomerular couplings. Comparison of homologous olfactory lineages across drosophilids reveals natural examples of fate changes from death to a functional neuron. Finally, we provide evidence that PCD contributes to evolutionary differences in carbon dioxide-sensing circuit formation in *Drosophila* and mosquitoes. These results reveal the remarkable potential of alterations in PCD patterning to evolve new neural pathways.

## INTRODUCTION

A fundamental way in which nervous systems evolve is through increases in the numbers of neurons (Han et al., 2015; Herculano-Houzel et al., 2015; Strausfeld et al., 2009). Additional sensory neurons can enable higher sensitivity to environmental signals or lead to functional diversification to support acquisition of novel detection abilities (Hansson and Stensmyr, 2011). Increases in central neuron number might underlie diverse enhancements in cognitive abilities (Fang and Yuste, 2017), such as parallel processing and memory storage.

The generation of more neurons could be achieved through greater production during development, by increasing the number and/or proliferation of neural precursor cells. This process appears to have contributed to neocortical expansion during primate evolution (Wilsch-Brauninger et al., 2016). Alternatively (or additionally), given the widespread occurrence of programmed cell death (PCD) during neural development (Dekkers et al., 2013; Yamaguchi and Miura, 2015), prevention of this process can potentially yield a pool of new neurons. Consistent with this idea, genetic blockage of PCD in mice or *D. melanogaster* results in the development of enlarged, albeit malformed, nervous systems (Kuida et al., 1998; Rogulja-Ortmann et al., 2007). In *C. elegans* lacking the CED-3 caspase (a key executioner of PCD), many of the surviving cells differentiate morphologically as neurons (White et al., 1991); moreover, one of these can partially compensate for the function of an experimentally-ablated sister pharyngeal neuron (Avery and Horvitz, 1987).

Here we examined the potential of PCD blockage to generate novel neural pathways in the *D. melanogaster* olfactory system. The principal olfactory organ in drosophilids, the third antennal segment (hereafter, antenna), is covered with ∼400 porous sensory hairs (sensilla) of morphologically-diverse classes (Figure 1A) (Shanbhag et al., 1999). An individual sensillum derives from a single sensory organ precursor (SOP) cell that is specified in the larval antennal imaginal disc (Endo et al., 2011; Rodrigues and Hummel, 2008). Each SOP gives rise to a short, fixed lineage of asymmetric cell divisions that produces eight terminal cells with distinct identities (Endo et al., 2007; Endo et al., 2011) (Figure 1B). Four adopt non-neural (“support cell”) fates, and are involved in the construction of the hair, amongst other roles. The other four cells can potentially differentiate as olfactory sensory neurons (OSNs), which each express a single (or, rarely, two) sensory receptor genes, develop ciliated dendrites that innervate the lumen of the sensillum hair, and project axons towards a specific glomerulus in the primary olfactory center (antennal lobe) in the brain. There are ∼20 sensillum classes, housing stereotyped combinations of OSNs (Table S1) (Benton et al., 2009; Couto et al., 2005; Grabe et al., 2016). Of these, only one class contains four neurons, with the others housing one, two or three OSNs. The “missing” neurons are removed by spatially-precise PCD ∼22-32 hours after puparium formation (APF) (Chai et al., 2019; Endo et al., 2007; Reddy et al., 1997; Sen et al., 2004), when OSN terminal fate is established (Jefferis and Hummel, 2006).

**Figure 1.**
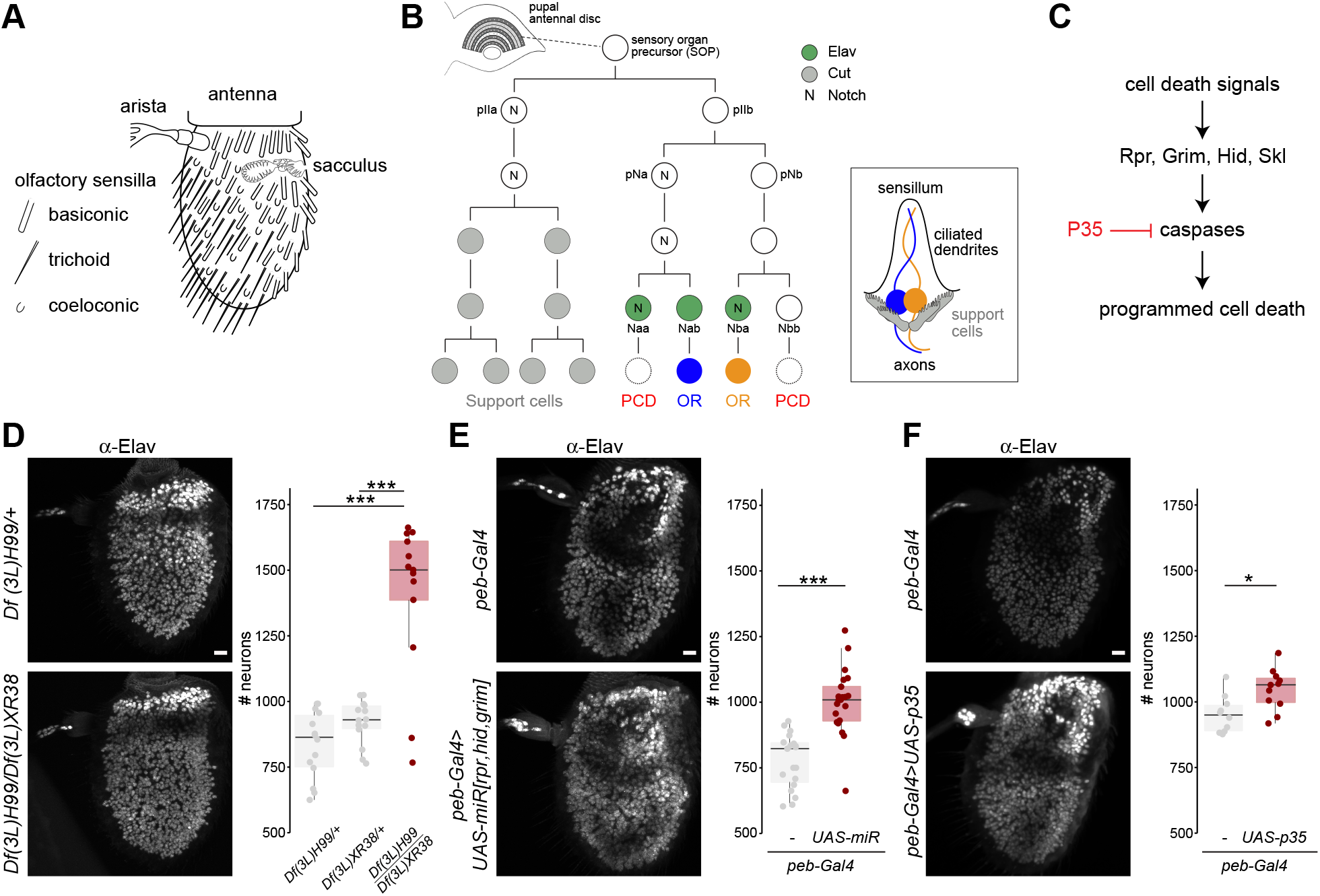
Inhibition of developmental programmed cell death results in increased neuron numbers in the antenna. **(A)** Schematic of the *Drosophila* third antennal segment showing the different sensory structures. **(B)** Schematic of the lineage of an antennal disc sensory organ precursor (SOP) cell giving rise to a sensillum containing two neurons (illustrated on the right). The expression of a subset of molecular markers is shown; Elav is expressed in only three of four neural precursors; one of these (Naa) as well as the Elav-negative cell (Nbb) are eliminated by PCD. The lineage is based upon data from (Chai et al., 2019; Endo et al., 2007; Endo et al., 2011). **(C)** Simplified schematic of the PCD pathway in *Drosophila*, highlighting the elements relevant for this study. Several intermediate steps between the pro-apoptotic proteins (Rpr, Grim, Hid and Skl) and the executioner caspases are not shown. **(D)** Elav expression in whole-mount antennae from control (*Df(3L)H99/+*; the wild-type chromosome here and in other genotypes was derived from a *w^1118^* parent) and PCD-deficient (*Df(3L)H99/Df(3L)XR38*) animals (here, mixed sexes were analyzed; in all other experiments, female flies were used, except where noted otherwise). Scale bar = 10 μm. Right: quantifications of antennal neuron numbers of the indicated genotypes, including an additional control genotype (*Df(3L)XR38)/+*) (n = 14, 14 and 13, respectively). *** indicates p = 0.0007216 for the comparison to *Df(3L)H99/+* and p = 0.0013224 for the comparison to *Df(3L)XR38/+* (Wilcoxon-sum rank test, corrected for multiple comparisons using a Bonferroni correction). In this and subsequent panels, individual data points are shown, overlaid with boxes indicating the median and first and third quartile of the data; whiskers showing the limits of the distribution. **(E)** Elav expression in whole-mount antennae from control (*peb-Gal4/+*) and PCD-blocked (*peb-Gal4/+;UAS-miR(grim,rpr,hid)/+*) animals. Scale bar = 10 µm. Right: quantifications of neuron numbers of these genotypes. *** indicates p = 0.0024×10^-4^ (t-test) (n = 19 and 21; control and PCD-blocked, respectively). **(F)** Elav expression in whole-mount antennae from control (*peb-Gal4/+*) and PCD-blocked (*peb-Gal4/+;UAS-p35/+*) animals. Scale bar = 10 µm. Right: quantifications of neuron numbers of these genotypes. * indicates p = 0.024 (t-test) (n = 10 and 11; control and PCD-blocked, respectively).

Here we demonstrate that prevention of cell death during the development of OSNs is sufficient to generate novel functional neurons that integrate within pre-existing olfactory circuits. Importantly, some of these undead neurons represent novel cell types as reflected by their reproducible receptor expression pattern, soma location, and axonal projections. Finally, we provide evidence for the evolutionary diversification of olfactory pathways through changes in deployment of PCD, both within drosophilid species, and between drosophilids and mosquitoes. We propose that cells normally fated to die represent an important evolutionary reserve for the generation of new neuron types and neural circuits.

## RESULTS AND DISCUSSION

### Inhibition of programmed cell death during development results in increased neuron number in the antenna

To block PCD during OSN development, we first used animals bearing deletions in the tandem cluster of pro-apoptotic genes (*head involution defective* (*hid*), *grim*, *reaper* (*rpr*) and *sickle* (*skl*)), which are critical for promoting developmentally-regulated PCD in diverse tissues (Figure 1C) (Lee et al., 2013; Pinto-Teixeira et al., 2016). Homozygous chromosomal deficiencies that span the entire cluster cause embryonic lethality. However, a trans-heterozygous combination (*Df(3L)H99/Df(3L)XR38*), which removes both copies of *rpr*, and one copy each of *hid*, *grim* and *skl*, allowed recovery of a few viable adults. Immunofluorescence on whole-mount antennae with an antibody against a neural nuclear marker, Elav, revealed a clear increase in the number of labeled cells in mutant animals compared to controls (Figure 1D), indicating that new neurons form when cell death is prevented.

PCD might be impaired in these mutant animals at any stage of olfactory system development, including during SOP specification in the antennal disc. To selectively block PCD in terminal OSN precursors (Figure 1B), we down-regulated expression of *hid*, *grim* and *rpr* simultaneously by transgenic RNAi from ∼18 h APF using the *pebbled-Gal4* (*peb-Gal4*) driver, which is broadly expressed in post-mitotic cells in these lineages (Sweeney et al., 2007). Blockage of OSN-specific PCD also led to a significant increase in Elav-positive cells (Figure 1E). The number of extra Elav-positive cells observed in this experiment (∼200-300, recognizing the limits of automated neuron counting in nuclei-dense antennal tissue (Figure S1)) is in line with estimates of the total number of potential “undead” neurons (∼300-400) (Table S1). We further confirmed the role of the canonical PCD pathway in the antenna through expression of the baculoviral caspase inhibitor P35 (Figure 1C) (Hay et al., 1994) with the same driver. *peb-Gal4>UAS-p35* (hereafter “PCD-blocked”) animals displayed higher numbers of Elav-positive cells compared to a *peb-Gal4* (control) genotype (Figure 1F), consistent with a caspase-dependent PCD pathway in this sensory organ.

### Undead olfactory sensory neurons are functional

To determine whether these additional Elav-positive cells are functional neurons, we performed single-sensillum electrophysiological recordings. We focused on one class of sensillum, antennal trichoid 1 (at1), which houses a single OSN in wild-type animals, due to PCD of the other three potential neurons in the lineage (Chai et al., 2019). This OSN expresses OR67d, a receptor for the pheromone *11-cis-*vaccenyl acetate (cVA) (Ha and Smith, 2006). at1 sensilla are easily recognized by their sparse basal (spontaneous) pattern of spikes of a single amplitude, and the robust train of spikes that occur upon presentation of cVA but not other odors (Figure 2A,D). In PCD-blocked animals, these sensilla often contain additional spikes of smaller amplitude (Figure 2A-C), suggesting the presence of one or more extra, active OSNs (spike amplitude is defined by the OSN not the receptor gene (Hallem et al., 2004)). Moreover, exposure to a blend of food-derived odors (which activate many different ORs (Hallem and Carlson, 2006)) led to responses of the undead neurons in about one-third of the tested sensilla (Figure 2D-E). The non-responding, but spiking, undead neurons might express a receptor activated by other stimuli. While this variability in responsiveness could reflect a stochastic fate of undead neurons, our analysis of the location and wiring of undead neurons presented below argues against this interpretation. We therefore suspect it is attributable to the PCD-blocking method not being fully efficient, resulting in random rescue of one or two different undead OSN types in distinct at1 sensilla. Regardless, these observations indicate that blocking PCD can lead to the development of functional OSNs.

**Figure 2.**
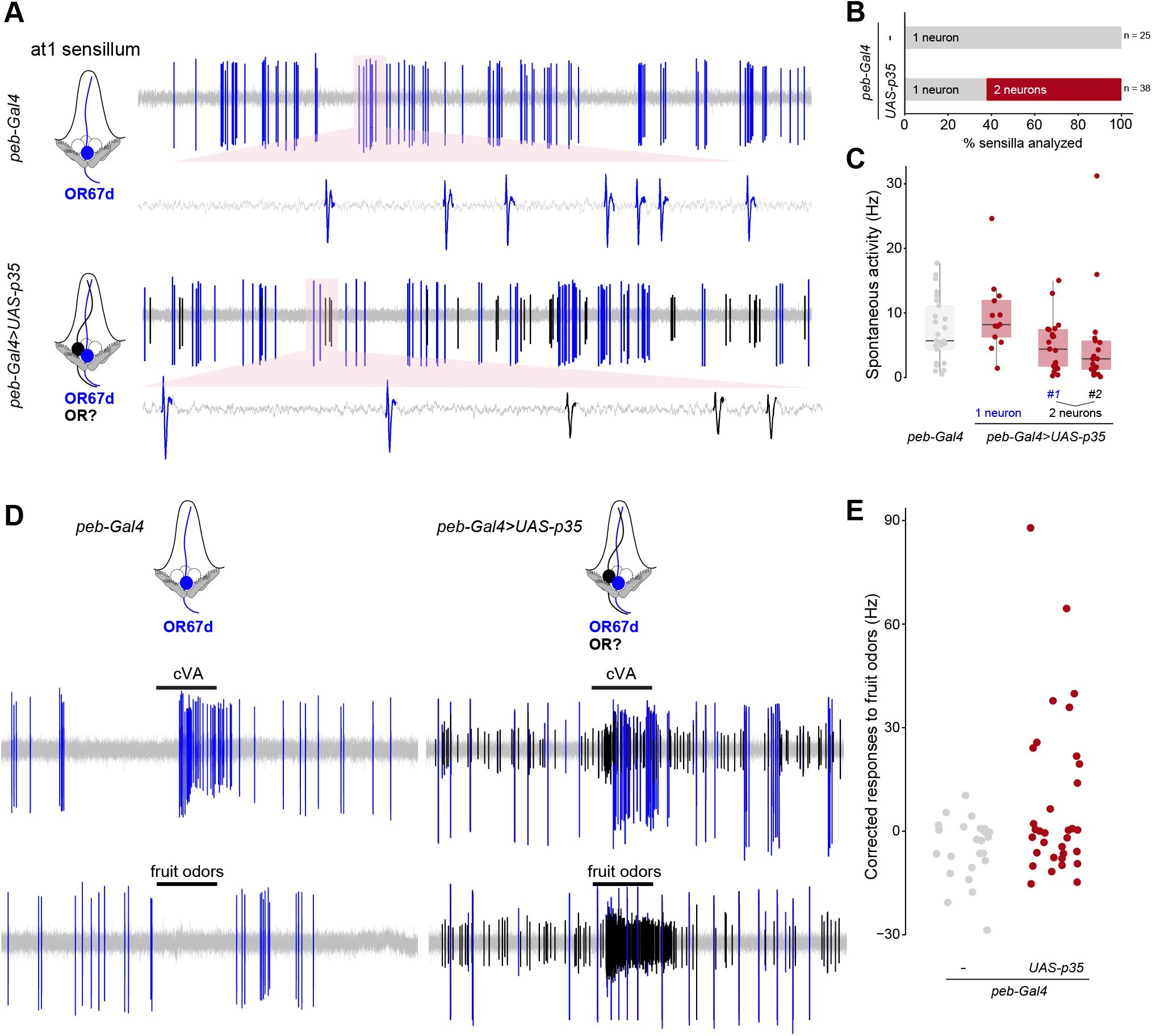
Undead olfactory sensory neurons are functional. **(A)** Representative extracellular electrophysiological traces of spontaneous activity from neurons in an at1 sensillum of control (*peb-Gal4/+*) and PCD-blocked (*peb-Gal4/+;UAS-p35/+*) animals. Automatically-detected spikes (see Methods) from the neuron expressing OR67d are shown in blue, and those of the additional, undead neuron(s) in black, as schematized in the cartoon on the left (cells fated to die are shown with dashed outlines). **(B)** Quantifications of the proportion of sensilla containing either one neuron (grey) or two (or more) neurons (red) in control (*peb-Gal4/+*) and PCD-blocked (*peb-Gal4/+;UAS-p35/+*) animals. **(C)** Quantifications of the spontaneous activity of the indicated neurons for the control and PCD-blocked genotypes. **(D)** Representative electrophysiological traces from at1 sensillum recordings in control (*peb-Gal4/+*) and PCD-blocked (*peb-Gal4/+;UAS-p35/+*) animals upon stimulation with a 0.5 s pulse (black horizontal bar) of the pheromone *11-cis*-vaccenyl acetate (cVA) (10^-2^ dilution (v/v) in paraffin oil) or a mix of fruit odors (butyl acetate, ethyl butyrate, *2*-heptanone, hexanol, isoamyl acetate, pentyl acetate; each odor at 10^-2^ dilution (v/v) in paraffin oil). Automatically-detected spikes from the neuron expressing OR67d are shown in blue, and those of the undead neuron(s) in black. **(E)** Quantifications of odor-evoked responses to fruit odors (see Methods) in control (*peb-Gal4/+*) and PCD-blocked (*peb-Gal4/+;UAS-p35/+*) animals (n = 25 and 34, respectively).

### Undead neurons express a subset of olfactory receptor genes

To identify the receptor genes expressed by undead OSNs, we performed comparative transcriptomics of whole antennae of control and PCD-blocked animals by RNA-sequencing. As a positive control, we first examined the changes in transcript levels of *grim*, *rpr*, *hid* and *skl*, reasoning that inhibition of PCD downstream in the pathway should lead to the presence of undead cells expressing mRNAs for these pro-apoptotic genes (Figure 1C). Indeed, three of these genes showed significantly higher expression levels in PCD-blocked antennae (Figure 3A and Table S2).

**Figure 3.**
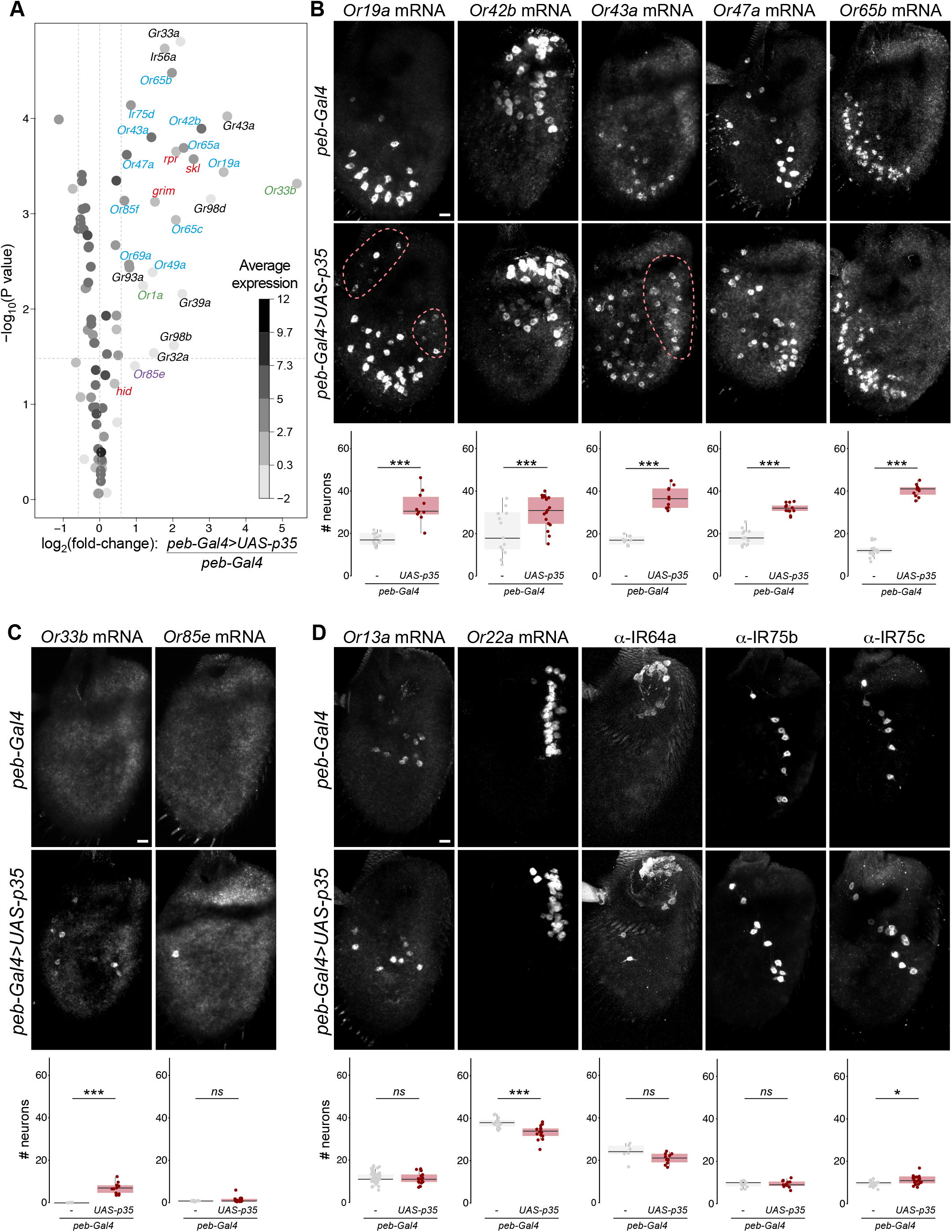
Undead neurons express a subset of olfactory receptor genes. **(A)** Gene expression differences between control and PCD-blocked antennae. The volcano plot shows the differential expression (on the x-axis) of *D. melanogaster Or*, *Ir* and *Gr* gene transcripts (each gene represented by a dot), as well as the four pro-apoptotic genes (*grim*, *rpr*, *hid* and *skl*; red labels), plotted against the statistical significance (on the y-axis). The mean expression level of individual genes across all samples is shown by the shading of the dot, as indicated by the grey scale on the right (units: log_2_(counts per million)). Only chemosensory genes showing a >1.5-fold increase in PCD-blocked antennae are labeled: blue labels indicate genes whose expression in the antenna has previously been demonstrated by RNA *in situ* hybridization; magenta and green labels indicate receptors normally only expressed in the adult maxillary palps and larval dorsal organ, respectively; black labels indicate receptors that are expressed in gustatory organs. The horizontal dashed line indicates a false discovery rate threshold of 5%. Data for all *Or*, *Ir*, *Gr* and pro-apoptotic genes are provided in Table S2. **(B)** Representative images of RNA FISH for the indicated *Or* genes in whole mount antennae of control (*peb-Gal4/+*) and PCD-blocked (*peb-Gal4/+;UAS-p35/+*) animals. Scale bar = 10 µm. Quantifications of neuron numbers are shown at the bottom. *** indicates Or19a p = 6.526×10^-5^ (t-test) (n = 17 and 10 (control and PCD-blocked, respectively)), Or42b p = 0.008486 (t-test) (n = 13 and 17), Or43a p = 5.888×10^-7^ (t-test) (n = 10 and 10), Or47a p = 3.088×10^-10^ (t-test) (n = 13 and 13), Or65b p = 2.2×10^-16^ (t-test) (n = 17 and 11) (see Figure S2A for additional examples). The pink dashed lines encircle cells in PCD-blocked antennae that express the corresponding *Or*s outside their usual spatial domain (see also Figure S3). **(C)** Representative images of RNA FISH for the indicated *Or* genes in whole mount antennae of control (*peb-Gal4/+*) and PCD-blocked (*peb-Gal4/+;UAS-p35/+*) animals. Scale bar = 10 µm. Quantifications of neuron numbers are shown at the bottom. *** indicates Or33a p = 1.812×10^-7^ (t-test) (n = 23 and 23), ns indicates Or85e p = 0.053 (Wilcoxon-sum rank test) (n = 12 and 22); although this is not statistically significant, we note we never detected any *Or85e* mRNA-positive neurons in control antennae, but frequently detected one (or more) labeled cells in PCD-blocked antennae. **(D)** Representative images of RNA FISH or immunofluorescence of the indicated olfactory receptors in whole mount antennae of control (*peb-Gal4/+*) and PCD-blocked (*peb-Gal4/+;UAS-p35/+*) animals (exceptionally, mixed sexes were analyzed for IR75b and IR75c immunofluorescence experiments). Scale bar = 10 µm. Quantifications of neuron numbers are shown at the bottom. ns, * and *** indicate Or13a p = 0.475 (t-test) (n = 46 and 17), Or22a p = 0.0002472 (t-test) (n = 15 and 15), Ir64a p = 0.128 (t-test) (n = 6 and 10), Ir75b p = 0.9246 (t-test) (n = 11 and 12), Ir75c p = 0.01 (t-test) (n = 19 and 18). Although there is not a statistically-significant increase in Ir64a neuron number, we reproducibly detect 1-2 extra IR64a-expressing cells outside the sacculus in PCD-blocked antennae (see also Figure S3).

We next queried the transcript levels for all chemosensory receptors, comprising *Odorant receptor* (*Or*), *Ionotropic receptor* (*Ir*) and *Gustatory receptor* (*Gr*) gene families (Tables S1-S2). Of the receptors previously detected in antennal neurons *in situ* (Benton et al., 2009; Couto et al., 2005; Fishilevich and Vosshall, 2005), we found that 10/36 *Or*s, 1/17 *Ir*s and 0/3 *Gr*s displayed a >1.5-fold increase in expression, suggesting that only subsets of these receptors are expressed in undead neurons (Figure 3A and Tables S2-S3).

To validate these transcriptomic data, we visualized the neuronal expression of several of the *Or*s *in situ*. Transcripts for all of those tested by RNA fluorescent *in situ* hybridization (FISH) were detected in more neurons in PCD-blocked antennae compared to controls (for *Or49a*, which we could not detect by RNA FISH, we used a validated *Or49a* promoter-CD8:GFP (*Or49a-*GFP) reporter (Couto et al., 2005)) (Figure 3B and Figure S2A). In some cases, these neurons were found only within the same region of the antenna as the endogenous OSNs (*e.g.,* Or42b, Or47a, Or65a, Or65b, Or85f) while in others (*e.g.*, Or19a, Or43a, Or49a) undead neurons were observed in novel, but reproducible, locations (Figure 3B, Figure S2A and Figure S3). The variance in the number of neurons expressing a particular receptor was not significantly different between control and PCD-blocked antennae (F-test for equality of variance, data not shown). Together, these observations suggest that undead neurons display consistent, rather than stochastic, receptor expression patterns.

Notably, many of the other receptors displaying increases in transcript levels normally act in other chemosensory organs, including one *Or* (*Or85e*) expressed in the maxillary palp (a distinct olfactory appendage of insects), two larval *Or*s (*Or1a* and *Or33b*), and seven *Gr*s, which function in various gustatory organs (Figure 3A and Tables S2-S3). *In situ* analysis confirmed the presence of transcripts for the palp-specific *Or85e* and the larval-specific *Or33b* in populations of undead neurons in PCD-blocked antennae (Figure 3C). These observations suggest that undead neurons can provide a cellular substrate to allow switching of receptor expression between sensory organs and/or life stages during evolution.

Beyond these cases, the RNA levels of the vast majority of receptor genes were either unchanged or slightly down-regulated in PCD-blocked antennae (Figure 3A and Table S2). Consistently, *in situ* analysis by FISH or immunofluorescence of these antennal receptors revealed only a very small increase (*e.g.*, Ir75c), no change (*e.g.*, Or13a, Or67d, Gr63a, Ir40a, Ir64a and Ir75b), or a decrease (*e.g.*, Or22a) in the size of the corresponding neuron populations (Figure 3D and Figure S2B). The latter, unexpected phenotype raises the possibility that undead neurons impact (directly or indirectly) the specification and/or survival of certain populations of neurons.

### Co-expressed receptor genes are over-represented in undead neurons

What properties characterize the small subset of receptors that are expressed in undead neurons? They are normally expressed in neurons housed in diverse sensillum types: basiconic (*e.g.*, *Or42b*), trichoid (*e.g.*, *Or65a*), intermediate (*e.g.*, *Or19a*), and coeloconic (*e.g.*, *Ir75d*). By contrast, most of these receptors (including 9/10 *Or*s) are expressed in OSNs derived from the Nba precursor cell; the remaining *Or* (*Or43a*) and the sole *Ir* (*Ir75d*) are expressed in Naa-derived OSNs (Figure 1B, Figure 4A and Table S3). This pattern suggests that undead neurons – which are largely Naa-derived (Figure 1B) – preserve gene-regulatory networks that are more similar to Nba cells than Nab cells, possibly reflecting the shared Notch activity in Nba and Naa precursors (Figure 1B) (Endo et al., 2011).

**Figure 4.**
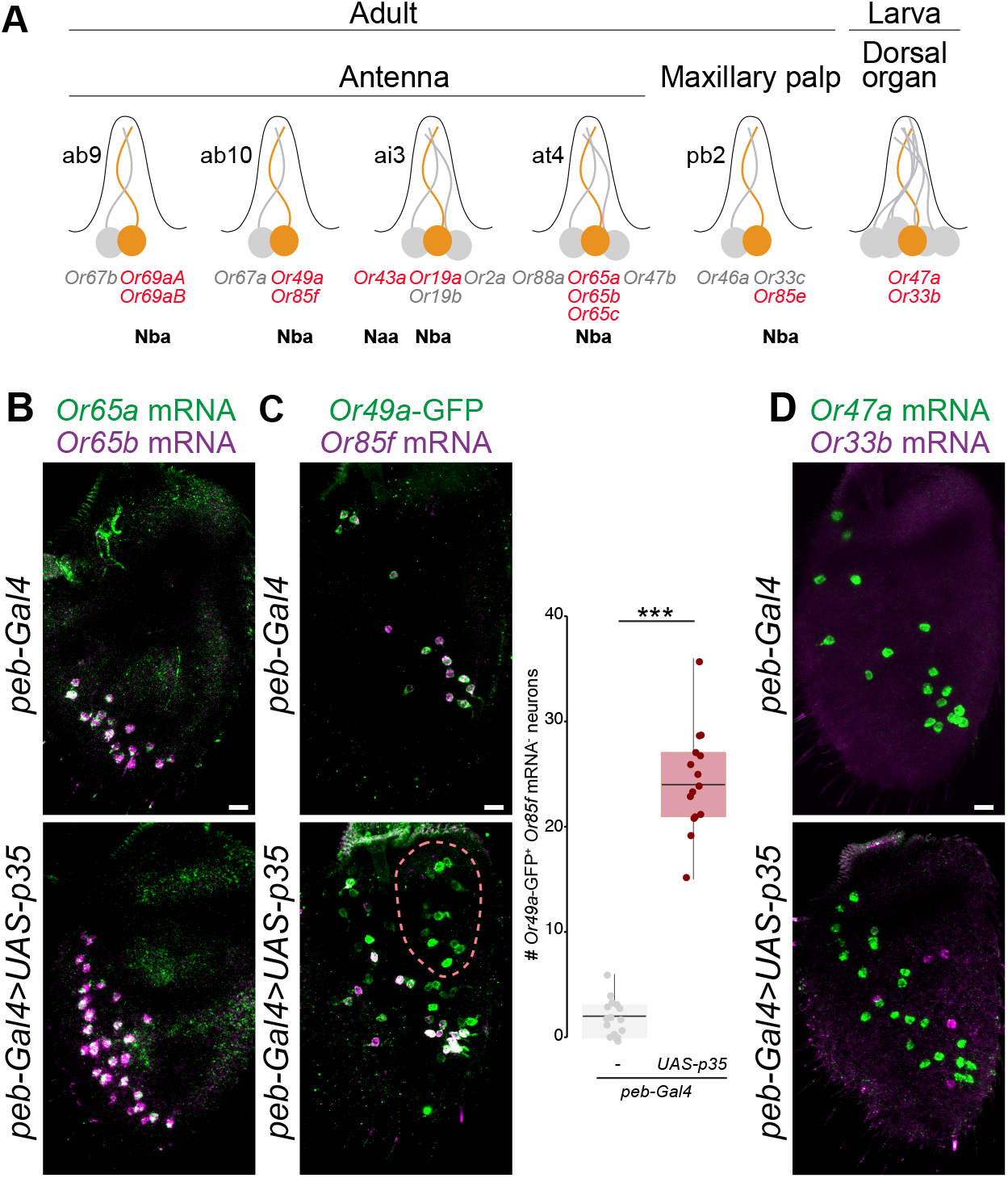
Co-expressed receptor genes are over-represented in undead neurons. **(A)** Schematic summarizing the normal olfactory organ/sensillum expression pattern of the subset of up-regulated *Or* genes that display co-expression in wild-type neurons (labeled in red; receptor genes showing no changes in transcript levels are labeled in grey). The neuronal precursor identity of these OSNs is shown below. **(B)** Representative images of combined RNA FISH for *Or65a* (green) and *Or65b* (magenta) in whole mount antennae of control (*peb-Gal4/+*) and PCD-blocked (*peb-Gal4/+;UAS-p35/+*) animals (n = 4 and 5, respectively), showing co-expression of these receptors in both endogenous and undead neurons. Scale bar = 10 µm. **(C)** Representative images of RNA FISH for *Or85f* and anti-GFP in whole mount antennae of control (*peb-Gal4/+;Or49a-GFP/+*) and PCD-blocked (*peb-Gal4/+;Or49a-GFP/UAS-p35*) animals. Scale bar = 10 μm. Co-expression quantifications are shown to the right. *** indicates *Or49a-GFP^+^/*Or85f mRNA^-^ population p = 5.48×10^-12^ (t-test) (n = 16 and 15 (control and PCD-blocked, respectively)). The pink dashed line encircles cells in the PCD-blocked antenna that express *Or49a*-GFP outside its usual spatial domain (see also Figure S3). We used an *Or49a-GFP* reporter due to our inability to reliably detect *Or49a* transcripts *in situ*; the higher number of *Or49a-CD8:GFP*-positive *Or85f* RNA-positive cells is not an artefact of the analysis method, as an *Or85f*-CD8:GFP reporter revealed a similarly limited increase in neuron number (Figure S4B). **(D)** Representative images of combined RNA FISH for *Or47a* (green) and *Or33b* (magenta) in whole mount antennae of control (*peb-Gal4/+*) and PCD-blocked (*peb-Gal4/+;UAS-p35/+*) animals. In control animals *Or33b* is co-expressed with *Or47a* in the larval dorsal organ, and is never detected in the antenna. In PCD-blocked antennae *Or33b*- and *Or47a*-expressing undead neurons are almost completely non-overlapping (only 4% of *Or33b*-positive undead OSNs weakly co-express *Or47a* (n = 79 cells from 10 antennae). Scale bar = 10 µm.

Finally, of the 13 *Or*s detected in undead neurons (including those from other olfactory organs), ten are normally co-expressed with other *Or* genes (Figure 4A and Table S3). This enrichment is striking given the rarity of receptor co-expression within this repertoire (Couto et al., 2005; Fishilevich and Vosshall, 2005). While some co-expressed receptors remain co-expressed in undead OSNs (*e.g.*, *Or65a* and *Or65b* (Figure 4B)), this is not always the case. For example, *Or19a*, but not the co-expressed *Or19b*, displays up-regulation by RNA-seq (Table S2). Similarly, while *Or49a-*GFP and *Or85f* are always co-expressed in control antennal OSNs, in PCD-blocked antennae we detected a population of undead neurons that expresses *Or49a*-GFP but not *Or85f* (Figure 4C). Finally, the larval receptor *Or33b* is co-expressed with *Or47a* at this life stage (Fishilevich et al., 2005), but in PCD-blocked antennae the novel *Or33b* undead neurons show only rare co-expression with *Or47a* (Figure 4D).

Why normally co-expressed receptors are over-represented in undead neurons remains unknown. We speculate that the chromatin state of these genes is more permissive for expression PCD-rescued cells. Regardless of its molecular basis, this phenomenon provides an intriguing link between receptor gene duplication – which is likely to be the first step in the evolution of a new olfactory channel (Ramdya and Benton, 2010) – and the formation of molecularly and cellularly distinct sensory pathways.

### Undead olfactory sensory neurons form novel receptor/glomerular couplings in the brain

We next investigated whether undead OSNs project their axons to the antennal lobe. We first broadly-labeled these neurons using an EGFP gene trap allele of *grim* (*grim^MI03811(EGFP)^*), in which the fluorophore should report on the expression pattern of this pro-apoptotic gene. In control animals, *grim^MI03811(EGFP)^* expression was detected only at background levels across the antenna; this is expected, as cells that induce Grim (and so EGFP) expression are fated to die (Figure 5A). By contrast, in PCD-blocked antennae, EGFP was present in many soma (Figure 5A), which presumably represent the undead neurons previously observed with Elav antibodies (Figure 1F). In the brains of these animals, we observed that EGFP-labeled processes innervate multiple glomeruli of the antennal lobe (Figure 5B), indicating that undead neurons can extend axons to the primary olfactory center. Antennal deafferentation experiments confirmed that the glomerular signals in PCD-blocked animals were entirely due to the contribution of OSNs (Figure 5B).

**Figure 5.**
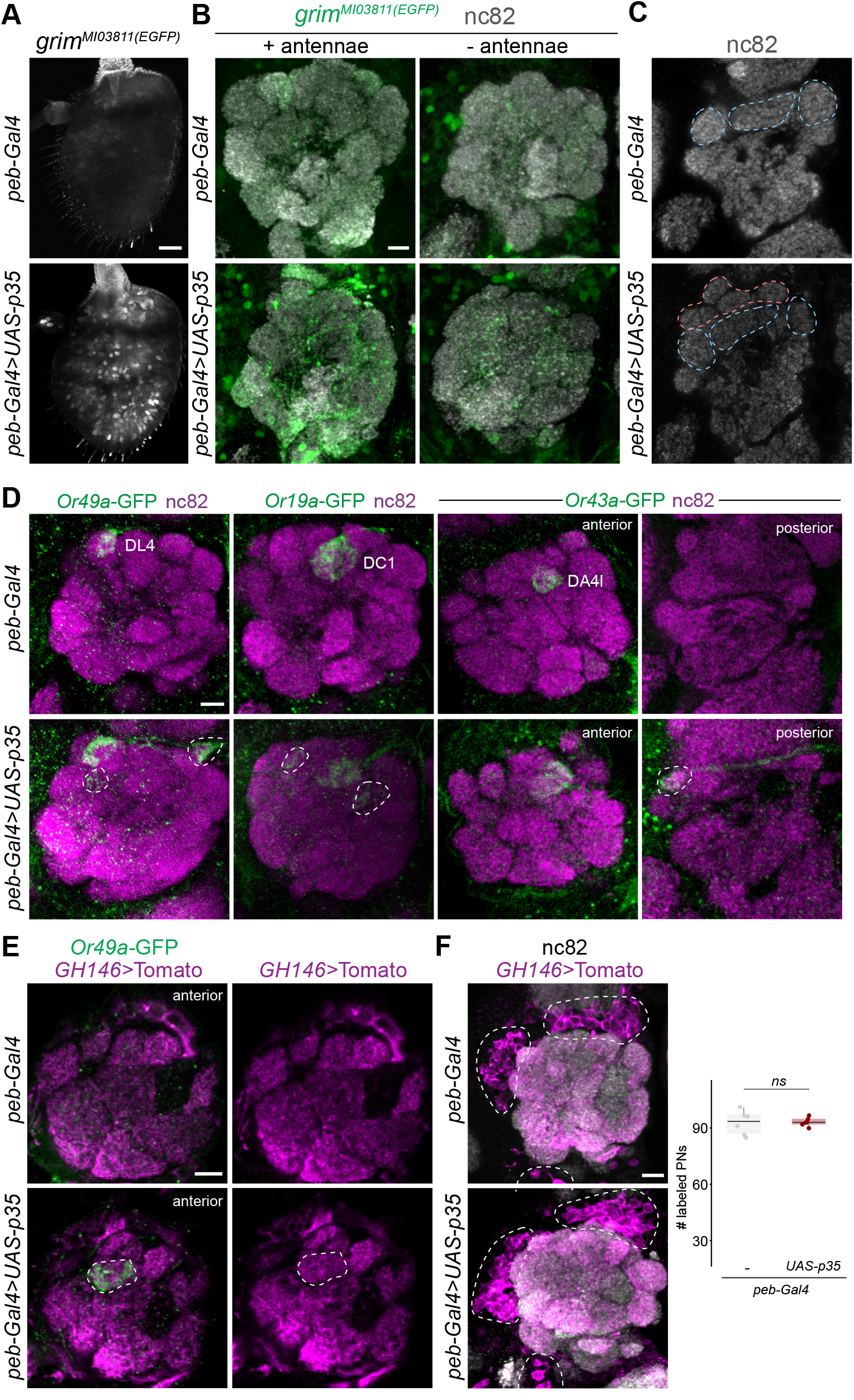
Undead olfactory sensory neurons form novel receptor/glomerular couplings in the brain. **(A)** Representative images of anti-GFP immunofluorescence in whole mount antennae of control (*peb-Gal4/+;;grim^MI03811(EGFP)^/+*) and PCD-blocked (*peb-Gal4/+; UAS-p35/+;grim^MI03811(EGFP)^/+*) animals. Blind scoring by two independent observers of antennae as belonging to control or PCD-blocked group was 100% accurate (n = 10 and 9 for control and PCD-blocked groups, respectively). Scale bar = 10 µm. **(B)** Representative images of combined anti-GFP and nc82 immunofluorescence in whole mount brains of control (*peb-Gal4/+;;grim^MI03811(EGFP)^/+*) and PCD-blocked (*peb-Gal4/+;UAS-p35/+;grim^MI03811(EGFP)^/+*) animals with intact (left) or excised antennae (right). Blind categorization of brains as belonging to the control (n = 9) or PCD-blocked (n = 12) set was 85% accurate (two independent observers). Scale bar = 10 µm. **(C)** Representative images of nc82 immunofluorescence in whole mount brains of control (*peb-Gal4/+;;grim^MI03811(EGFP)^/+*) and PCD-blocked (*peb-Gal4/+;UAS-p35/+;grim^MI03811(EGFP)^/+*) animals. Equivalent glomeruli found in antennal lobes of control and PCD-blocked animals are labeled for reference with dashed blue lines; potential novel glomerular structures (or displaced original glomeruli) in PCD-blocked animals are shown with dashed pink lines. Further planes of the same brains are shown in Figure S5. Blind categorization of brains as belonging to the control (n = 11) or PCD-blocked (n = 13) set (visualizing only the nc82 channel of confocal stacks of animals containing a diversity of fluorescent reporters in the genetic background; see genotypes below) was 85% accurate (two independent observers). Scale bar = 10 µm. **(D)** Representative images of combined anti-GFP and nc82 immunofluorescence in whole mount brains of control (*peb-Gal4/+;Or49a-GFP/Or49a-GFP, peb-Gal4/+;;Or19a-GFP/+, peb-Gal4/+;Or43a-GFP/+*) and PCD-blocked (*peb-Gal4/+;Or49a-GFP/Or49a-GFP,UAS-p35, peb-Gal4/+;UAS-p35/+;Or19a-GFP/+*, *peb-Gal4/+;Or43a-GFP/UAS-p35*) animals. Blind categorization of brains as belonging to the control or PCD-blocked set (visualizing only the GFP channel of confocal stacks and based exclusively on additional glomerular innervation patterns) was, for each reporter, respectively: *Or49a*-GFP: 85% (n = 10 and 10 for control and PCD-blocked); *Or19a*-GFP: 89.5% (n = 20 and n = 18); *Or43a*-GFP: 93.9% (n = 19 and n = 15). Scale bar = 10 µm. Additional representative images are provided in Figure S6. **(E)** Representative images of combined anti-GFP, anti-RFP and nc82 immunofluorescence in whole mount brains of control (*peb-Gal4/+;Or49a-GFP/Or49a-GFP;GH146-QF,QUAS-Tomato/+*) and PCD-blocked (*peb-Gal4/+;Or49a-GFP/Or49a-GFP,UAS-p35;GH146-QF,QUAS-Tomato/+*) animals. The dashed line encircles the novel *Or49-*GFP labeled glomerulus (*i.e.*, not the endogenous DL4 glomerulus, which is not visible in this anterior plane of the antennal lobe). Scale bar = 10 µm. **(F)** Representative images of PN soma (bounded by the dashed lines) labeled by GH146>Tomato in whole mount brains of control (*peb-Gal4/+;Or49a-GFP/Or49a-GFP;GH146-QF,QUAS-Tomato/+*) and PCD-blocked (*peb-Gal4/+;Or49a-GFP/Or49a-GFP,UAS-p35;GH146-QF,QUAS-Tomato/+*) animals. Scale bar = 10 µm. Quantifications of labeled PN numbers are shown to the right. *ns* indicates p = 0.819 (t-test) (n = 6 and 5, control and PCD-blocked, respectively).

Analysis of the overall architecture of the antennal lobe in control and PCD-blocked animals, as visualized with the synaptic marker nc82 (Bruchpilot) (Wagh et al., 2006), revealed significant morphological differences (Figure 5C and Figure S5; blind classification of genotype by two experimenters 85.3% accurate, n = 34), including less distinct boundaries between certain glomeruli and putatively novel regions of neuropil. To test whether these differences reflect the innervation patterns of populations of undead OSNs, we examined the projections of the neurons expressing reporters for several of the *Or* populations that increase in size in PCD-blocked antennae: *Or49a*-GFP, *Or19a-*GFP and *Or43a*-GFP (Figure 5D, Figure S4A and Figure S6). In control animals *Or49a*-GFP neurons project to a single glomerulus, DL4, as previously described (Couto et al., 2005). In PCD-blocked animals, labeled axons projected to DL4, as well as to a second, more anterior, glomerulus-like structure, and occasionally to a more medial location (Figure 5D and Figure S6A; blind classification of genotype 85% accurate, n = 20). These presumably correspond to the wild-type Or49a neuron population and the undead neurons that express this reporter (but not *Or85f* (Figure 4C)), respectively. Similarly, endogenous neurons expressing *Or19a*-GFP project to a single glomerulus, DC1, whereas undead *Or19a*-GFP*-*expressing neurons target two additional regions of neuropil (Figure 5D and Figure S6A; blind classification of genotype 89% accurate, n = 38). Finally, endogenous *Or43a*-GFP-expressing OSNs project to DA4l, while the undead neurons expressing this reporter innervate a distinct posterolateral glomerulus (Figure 5D and Figure S6A; blind classification of genotype 94% accurate, n = 34).

The reproducibility of projection patterns of undead neurons of a given type – together with the characteristic location of undead neuron soma in the antenna (Figure 3B and Figure S3) – is consistent with undead neurons adopting a more-constrained, rather than completely random, developmental fate. The global changes in glomerular boundaries make it difficult to distinguish whether the axons of undead neurons in these three cases target novel glomeruli, or partially/completely overlap with pre-existing glomeruli (*i.e.*, those formed by other populations of endogenous neurons). Nevertheless, the differences in projection patterns of undead and endogenous neurons that express the same receptor highlights that undead neurons may allow novel coupling between receptor expression and glomerular target, an essential step during the evolution of new olfactory pathways.

We next asked whether undead OSN axons can potentially synapse with second-order projection neurons (PNs), which carry olfactory signals to higher brain regions (Masse et al., 2009). We combined the *Or*-GFP reporters with a genetic driver for many PNs (*GH146-QF>QUAS-Tomato*) in control and antennal PCD-blocked flies. In the novel *Or49a*-GFP-labeled glomerulus, *GH146*-labeled processes were detected (Figure 5E). The undead neuron glomeruli innervated by *Or19a*-GFP and *Or43a*-GFP did not overlap with *GH146*-positive PNs (Figure S6B) but we suspect that this absence is due to incomplete coverage of PNs by the *GH146*-QF driver (we note that it does not label the endogenous *Or19a/*DC1 glomerulus (Figure S6B)). Importantly, nc82 immunoreactivity is present in the undead neuron glomeruli of all three OSN classes (Figure 5D), implying the formation of synapses between these sensory neurons and central circuit elements. Novel connectivity of undead OSNs does not result from the production of additional PNs (Figure 5F). This result suggests that there is no mechanism to match OSN and PN numbers, and that the observed innervation derives from local recruitment of PN dendrites in the antennal lobe during synapse formation. How new PN classes dedicated to novel OSN populations evolve remains unknown.

### Natural examples of evolved differences in olfactory sensory neuron numbers

Our demonstration that inhibition of PCD is sufficient to allow the development of new functional OSN populations that integrate into the olfactory circuitry is consistent with the hypothesis that modulation of cell death patterns during evolution can be a mechanism to create (or, conversely, remove) olfactory channels. While the variation in OSN number per sensilla within *D. melanogaster* (Table S1) implies that different SOP lineages have distinct regulation of PCD, we wondered whether we could identify examples of divergent deployment of PCD across shorter evolutionary timescales by comparing numbers of neurons in homologous olfactory sensilla in different drosophilids. Previous cross-species analyses suggested that this is likely to be relatively rare, as no differences in neuron numbers were reported in at least a subset of basiconic and coeloconic sensilla in a limited range of drosophilids (even though receptor tuning properties can vary substantially) (*e.g.*, (de Bruyne et al., 2010; Prieto-Godino et al., 2017; Stensmyr et al., 2003)).

We therefore performed a broader electrophysiological screening in 26 drosophilid species, focusing on the at1 sensillum class for ease of neuronal spike amplitude sorting. While at1 sensilla of most species house a single cVA-responsive neuron (Figure 6A-B), similar to *D. melanogaster*, we identified nine species in which this sensillum class houses two neurons of distinct spike amplitudes (Figure 6C), only one of which is cVA-responsive. The lack of genomic data for these species (except *D. mojavensis wrigleyi*) currently precludes further molecular analysis, but we assume that the cVA-responsive at1 neuron expresses an OR67d ortholog and the partner neuron a distinct receptor of still-unknown sensory specificity. We have unfortunately not been able to determine the receptor(s) expressed in undead neurons in *D. melanogaster* at1 sensilla (data not shown), but given the phylogenetic distance between these drosophilids, it is possible – if not likely – that these natural additional at1 neurons express receptors not even present in *D. melanogaster*.

**Figure 6.**
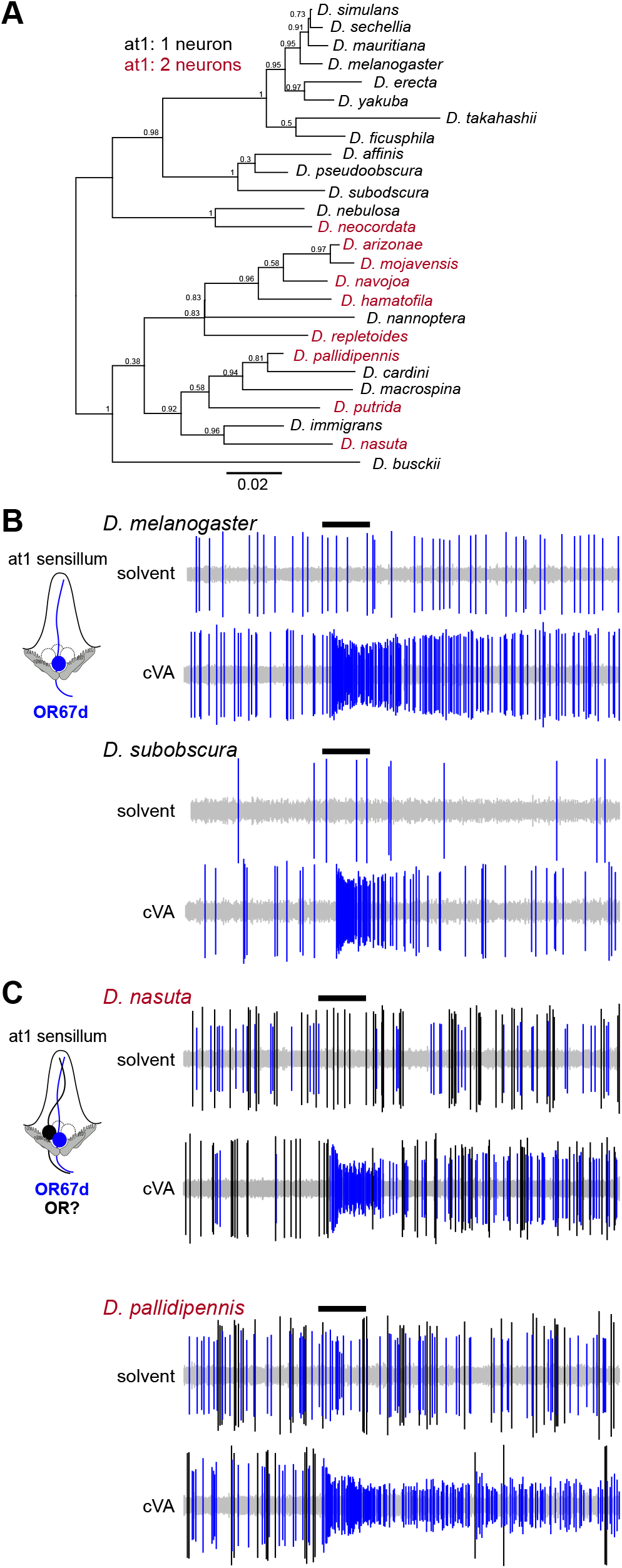
Examples of naturally occurring extra neurons in at1 sensilla. **(A)** Phylogeny of 26 drosophilid species, representing the majority of the *Drosophila* genus subgroups, based on the protein sequences of housekeeping loci (see Methods). Species names are colored to reflect the presence of one or two neurons in at1 sensilla. Numbers next to the tree nodes indicate the support values. The scale bar for branch length represents the number of substitutions per site. **(B)** Representative traces of extracellular recordings of neuronal responses to a 0.5 s pulse (black horizontal bar) of solvent (dichloromethane) or cVA in *D. melanogaster* (n = 5) and *D. subobscura* (n = 5). A single cVA-responsive neuron (known or assumed to express OR67d orthologs) is detected (blue spikes), as schematized in the cartoon on the left. **(C)** Representative traces of extracellular recordings of neuronal responses to a 0.5 s pulse (black horizontal bar) of solvent (dichloromethane) or cVA in *D. nasuta* (n = 5) and *D. pallidipennis* (n = 5). Two spike amplitudes are detected: a cVA-responsive neuron (assumed to express OR67d orthologs) (blue spikes) and second neuron with a larger spike amplitude, which does not respond to cVA (black spikes), as schematized in the cartoon on the left.

Although we cannot exclude that the extra at1 neuron in these nine species is due to an extra cell division, it would be an unprecedented property of an SOP lineage to have only one of the four terminal cells undergo an additional division. The most plausible explanation for the at1 phenotype in these species is that it reflects a change in fate from PCD to a functional OSN to permit formation of a novel olfactory channel. Mapping the species whose at1 sensilla house more than one OSN onto a phylogenetic tree reveals that the acquisition of an additional neuron (*i.e.*, a potential change in PCD patterning) has occurred independently multiple times during the evolution of the drosophilid clade (Figure 6A). This observation suggests that the diversification in sensilla development has a relatively simple – and potentially common – genetic basis.

### Programmed cell death can explain an evolutionary difference in carbon dioxide-sensing neuron formation in drosophilids and mosquitoes

Differences in neuron numbers within homologous sensilla have been described in the maxillary palp of more divergent dipteran species. In *D. melanogaster* (and other drosophilids), palp sensilla each house two *Or*-expressing OSNs (de Bruyne et al., 1999; Dweck et al., 2016; Singh and Nayak, 1985). By contrast, in mosquitoes, these sensilla house three neurons, comprising two *Or*-expressing OSNs and the carbon dioxide (CO_2_)-sensing neuron (expressing CO_2_ receptor subunits encoded by *Gr* genes) (Figure 7A) (Lu et al., 2007; McMeniman et al., 2014). In drosophilids, CO_2_-sensing neurons (expressing orthologous *Gr* genes) are confined to the antenna (Figure 7A) (Jones et al., 2007; Kwon et al., 2007).

**Figure 7.**
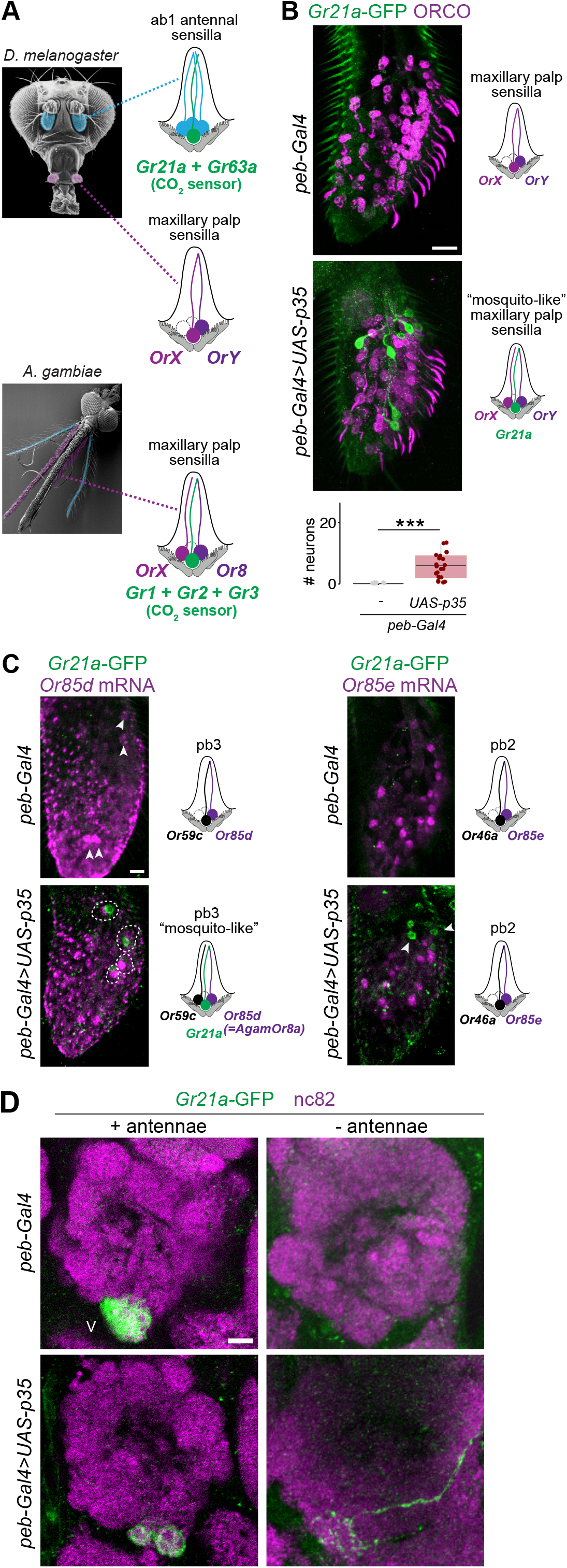
Programmed cell death can explain an evolutionary difference in carbon dioxide-sensing neuron formation in *D. melanogaster* and mosquitoes. **(A)** Left: SEM images of *D. melanogaster* and *A. gambiae* heads indicating the two olfactory organs: antennae (blue) and maxillary palps (magenta) (adapted from (Ramdya and Benton, 2010)). Right: schematic illustrating that in *D. melanogaster* CO_2_-sensing neurons (green) are located in antennal ab1 sensilla and that maxillary palp sensilla house only two OR-expressing neurons. By contrast, in *A. gambiae* (and other mosquitoes), CO_2_-sensing neurons are located in the maxillary palp, housed together with two OR-expressing neurons. **(B)** Representative images of combined anti-ORCO (the OR co-receptor, which labels all OR neurons (Larsson et al., 2004)) and anti-GFP immunofluorescence in whole mount maxillary palps of control (*peb-Gal4/+;Gr21a-GFP/+*) and PCD-blocked (*peb-Gal4/+;Gr21a-GFP/UAS-p35*) *D. melanogaster*. Scale bar = 10 μm. Quantifications of *Gr21a-*GFP-positive neuron numbers are shown at the bottom; *** indicates p = 6.7×10^-9^ (Wilcoxon sum-rank test) (n = 22 and n = 21 (control and PCD-blocked, respectively)). Schematic of inferred maxillary palp sensilla configuration in each genotype is illustrated on the right. **(C)** Representative images of combined RNA FISH for the indicated *Or* genes (magenta) and anti-GFP immunofluorescence (green) in whole mount maxillary palps of control (*peb-Gal4/+;Gr21a-GFP/+*) and PCD-blocked (*peb-Gal4/+;Gr21a-GFP/UAS-p35*) *D. melanogaster*. GFP-positive neurons pair with Or85d OSNs in pb3 sensilla, but not Or85e OSNs in pb2 sensilla. **(D)** Representative images of combined anti-GFP and nc82 immunofluorescence in whole mount brains of control (*peb-Gal4/+;Gr21a-GFP/+*) and PCD-blocked (*peb-Gal4/+;Gr21a-GFP/UAS-p35*) animals with intact (left) or excised antennae (right). Blind categorization of brains from excised antennae animals as belonging to the control or PCD-blocked set (visualizing only the GFP channel of confocal stacks and based exclusively on the presence of GFP-labeled axons) was 94.4% accurate (n = 19 and n = 17 for control and PCD-blocked, respectively). Scale bar = 10 μm.

To investigate whether PCD can account for differences in palp sensilla neuron organization, we examined a reporter of the CO_2_ receptor subunit GR21a in control and PCD-blocked *D. melanogaster*. *Gr21a*-GFP is expressed in the antenna but not the palps of control animals (Figure 7B and data not shown), consistent with previous observations (Jones et al., 2007; Kwon et al., 2007); remarkably, in PCD-blocked palps, this reporter labels several neurons (Figure 7B). Co-stainings with RNA probes for maxillary palp *Or*s revealed that these *Gr21a*-GFP-positive neurons are paired specifically with OSNs expressing *Or85d* in the pb3 sensillum class (Figure 7C). This is notable because *Or85d* is the *D. melanogaster* receptor that is most closely related to mosquito *Or8* ((Matthews et al., 2018) and data not shown), which is one of the two *Or*s expressed in mosquito palp sensilla (Figure 7A). Thus, prevention of PCD creates a sensillum in *D. melanogaster* maxillary palps that molecularly resembles sensilla organization in the homologous mosquito sensory organ.

Finally, we examined where these palp *Gr21a*-GFP-expressing undead neurons project. In control animals, antennal *Gr21a-*GFP neurons project unilaterally to the V glomerulus (Figure 7D) (as described (Jones et al., 2007; Kwon et al., 2007)). In PCD-blocked animals, *Gr21a-*GFP neurons (comprising both antennal and undead palp neuron populations) also converge on the V glomerulus. However, we could detect commissural projections and/or ectopic sites of innervation adjacent to this glomerulus (Figure 7D). Antennal deafferentation allowed us to selectively visualize *Gr21a-*GFP undead palp neurons: in controls, as expected, no innervations are labeled, while in PCD-blocked animals GFP-positive palp axons project bilaterally, and run amongst and occasionally branch within the medial glomeruli of the antennal lobe before terminating within (or very near) the V glomerulus (Figure 7D and Figure S8). The projection pattern of these undead neurons is partly reminiscent of mosquito CO_2_-sensing palp neurons, which project bilaterally to medial glomeruli (Ghaninia et al., 2007; Riabinina et al., 2016).

These results indicate that changes in PCD patterning can explain, in part, an evolutionary difference between drosophilids and mosquitoes for this important olfactory pathway. Additional genetic modifications must have occurred, for example, to promote high-level expression of CO_2_-receptor genes (which we have not yet been to able detect by RNA FISH) and to fully distinguish the axonal projection patterns of these neurons. Such developmental differences may be controlled – directly or indirectly – by the transcription factor Prospero and/or the microRNA, *mir-279*, whose loss in *D. melanogaster* leads to formation of ectopic palp CO_2_ neurons (Cayirlioglu et al., 2008; Hartl et al., 2011).

## CONCLUDING REMARKS

We have shown that a simple genetic manipulation preventing PCD is sufficient to allow development of functional OSNs that integrate into the extant olfactory circuitry of *D. melanogaster*, and provide evidence that species-specific deployment of PCD contributes to differences in olfactory pathway organization across drosophilids and mosquitoes.

A future challenge will be to understand the natural mechanisms controlling PCD in OSN lineages in order to address how this process is activated or suppressed during evolution to selectively eliminate or create OSNs. Our RNA-seq dataset provides an entry-point to answer this question by identifying candidate genes expressed highly in cells normally fated to die, similar to the pro-apoptotic factors. Given the timing and precise stereotypy of PCD within the olfactory lineages (Chai et al., 2019; Endo et al., 2007; Endo et al., 2011), it is also possible that PCD is induced through similar mechanisms to those that define the fate of OSNs that survive and express specific receptor genes (Barish and Volkan, 2015).

One key observation of our study is that undead neuron populations do not necessarily exhibit functional or anatomical properties that match those of existing OSNs, as exemplified by their expression of receptor genes not normally activated in antennal neurons, and by the existence of new receptor/glomerular couplings. These traits presumably reflect properties of undead OSNs’ inherent (though normally “hidden”) gene regulatory networks, and reveal the evolutionary potential of cells fated to die to differentiate as neurons with novel functions and wiring patterns. Future work should reveal how alterations in PCD patterning combine with other genetic changes (e.g., to refine receptor expression and neuronal projections) to form new, precisely-segregated olfactory pathways. Given the widespread occurrence of PCD throughout the nervous systems of insects and other animals (Dekkers et al., 2013; Yamaguchi and Miura, 2015), the contribution of undead neurons to neural circuit evolution is likely to extend to many different brain regions in diverse species (Pop et al., 2019).

## Acknowledgements

We are grateful to I. Alali and M. Erdogmus for technical assistance, Darren Williams and Sînziana Pop for sharing flies and discussions, Raquel Álvarez Ocaña for sharing information on neuron numbers, Mattias Alenius, Liqun Luo, Leslie Vosshall, the Bloomington *Drosophila* Stock Center (NIH P40OD018537), and the Developmental Studies Hybridoma Bank (NICHD of the NIH, University of Iowa) for reagents. We thank Roman Arguello and members of the Benton laboratory for comments on the manuscript. L.L.P.-G. was supported by a FEBS Long-Term Fellowship and an ERC Starting Independent Researcher Grant (802531). M.A.K., B.S.H., and M.K. are supported by the Max Planck Society. Research in R.B.’s laboratory is supported by the University of Lausanne, an ERC Consolidator Grant (615094) and the Swiss National Science Foundation.

## Author contributions

L.L.P.-G. and R.B. conceived the project. All authors contributed to experimental design, analysis and interpretation of results. Experimental contributions were as follows: L.L.P.-G. (Figures 2, 3D, 5A, S1); A.F.S. (Figures 1D-F, 3D, 4C, S1, 5B-C,E,F, S2A, S3, S4B, S5), M.A.K. (Figures 6, S7), S.C. (Figures 3, 4B-D, 7B-C, S2, S3, S4A), R.B. (Figures 5D, 7D, S6, S8). K.B. and S.P. performed RNA-seq data analysis. R.B., L.L.P.-G. and A.F.S. wrote the paper with input from all other authors.

## METHODS

### Drosophila culture

Flies were maintained at 25°C in 12 h light:12 h dark conditions, except where noted. *D. melanogaster* strains were cultured on a standard cornmeal diet; other drosophilid species were grown on food sources as indicated in the Key Resources Table (for recipes: http://blogs.cornell.edu/drosophila/recipes). Published mutant and transgenic *D. melanogaster* are described in the Key Resources Table. *Df(3L)H99/Df(3L)XR38* (and their controls) were cultured at 22°C to increase the recovery of adult offspring of the desired genotype. For most histological experiments, only female flies were analyzed, to avoid confounding variation due to known sexual dimorphisms (Grabe et al., 2016). Mixed sexes were used for *Df(3L)H99/Df(3L)XR38* flies in Figure 1D due to the limitation in the recovery of this genotype, as well as for anti-IR75b and anti-IR75c immunofluorescence in Figure 3D (there is no sexual dimorphism in the numbers of these OSNs). For histological experiments, flies were 1-12 day old. Animals subjected to antennal deafferentation (and control intact flies) were left for 10 days post-surgery to permit degeneration of OSN axons. For the experiments in Figure 6 and Figure S7, all experiments were carried out with 8-15 day old, mated female flies.

### Histology

Whole mount antennal and maxillary palp immunofluorescence and RNA fluorescent *in situ* hybridization were performed essentially as described (Saina and Benton, 2013). Whole mount brain immunofluorescence was performed essentially as described (Sanchez-Alcaniz et al., 2017). Primary and secondary antibodies are listed in the Key Resources Table; concentrations used are listed in Table S4. Sources and/or construction details of templates for RNA probes are provided in the Key Resources Table. Imaging was performed on a Zeiss confocal microscope LSM710 or LSM880 using a 40x oil immersion objective.

### Electrophysiology

Single-sensillum recordings were performed and analyzed essentially as described (Benton and Dahanukar, 2011; Olsson and Hansson, 2013). at1 sensilla were identified based upon their morphology and characteristic distal distribution on the antenna; they could also be clearly distinguished from the only other trichoid sensillum class, at4, which houses three OSNs (Figure S7 and data not shown). Chemical stimuli and solvents are described in the Key Resources Table. For the experiments in Figure 2, neuron activity was recorded for 10 s, starting 3 s before a stimulation period of 0.5 s. For the experiments in Figure 6 and Figure S7, neuron activity was recorded for 6 s, starting 2 s before a stimulation period of 0.5 s.

### RNA-sequencing

Antennal RNA was extracted from three biological replicates of control (*peb-Gal4/+;Or49a-GFP/+*) and PCD-blocked (*peb-Gal4/+;Or49a-GFP/UAS-p35*) animals. (The increased numbers of neurons labeled by *Or49a-*GFP was noted in preliminary studies and we therefore incorporated this transgene into the genotypes used in these experiments as an internal control; see below). For each pair of biological replicates, ∼200 animals were grown under identical conditions and RNA was extracted in parallel using 2-5 day old flies, as described (Chai et al., 2019). RNA quality was assessed on a Fragment Analyzer (Advanced Analytical Technologies, Inc.); all RNAs had an RQN of 9.8-10. From 100 ng total RNA, mRNA was isolated with the NEBNext Poly(A) mRNA Magnetic Isolation Module. RNA-seq libraries were prepared from the mRNA using the NEBNext Ultra II Directional RNA Library Prep Kit for Illumina (New England Biolabs). Cluster generation was performed with the resulting libraries using the Illumina TruSeq PE Cluster Kit v4 reagents and sequenced on the Illumina HiSeq 2500 using TruSeq SBS Kit v4 reagents (Illumina). Sequencing data were demultiplexed using the bcl2fastq Conversion Software (version 2.20, Illumina).

### Phylogenetics

Phylogenetic analysis of drosophilid species was conducted using six housekeeping proteins, encompassing two nuclear loci (*Adh* and Xdh) and four mitochondrial loci (*COI*, *COII*, *COIII* and *ND2*). Available amino acid sequences from Uniprot (https://www.uniprot.org, accession numbers are listed in Table S5) of each species were concatenated in Geneious (v11.0.5). A multiple sequence alignment of 2939 positions was generated using the MAFFT (v7.309) tool with E-INS-I parameters and scoring matrix 200 PAM / K=2 (Katoh and Standley, 2013). The final tree was reconstructed using a maximum likelihood approach with the GTR+G+I model of nucleotide substitution and 1000 rate categories of sites in FastTree (v2.1.5). The tree was visualized and processed in Geneious (v11.0.5).

## QUANTIFICATION AND STATISTICAL ANALYSIS

### Image analysis

For automated counting of Elav-positive cell bodies, confocal stacks were imported into Fiji (Schindelin et al., 2012) and passed through a median 3D filter of radius 1 in all dimensions. Images were subsequently thresholded using the 3D iterative thresholding plug-in (Ollion et al., 2013), and cells automatically counted using the 3D object counter.

For manual counting of OSN numbers expressing specific olfactory receptor genes, confocal stacks were imported into Fiji (Schindelin et al., 2012), and cell counting was performed manually using the ‘‘Cell counter’’ plug-in of ImageJ.

Analyses of OSN numbers expressing specific olfactory receptor genes, and morphological differences of the antennal lobes of control and PCD-blocked animals were performed by experimenters blind to the genotype, using RandomNames.bat (https://github.com/DavidOVM/File-Name-Randomizer/blob/master/RandomNames.bat) to encode image names.

### Electrophysiological analysis

Traces were analyzed by sorting spike amplitudes in AutoSpike (Syntech); representative traces presented in the figures were further processed in Adobe Illustrator CS (Adobe systems, San Jose, CA). Spontaneous neuron activity was quantified by counting spontaneous spikes in a 10 s recording window. Stimulus-evoked activity was quantified by counting spikes in a 0.5 s window during odor stimulation, and then subtracting this count from a 0.5 s recording window prior to stimulation. For the solvent-corrected quantifications in Figure 2D-E, the responses to solvent (paraffin oil) were subtracted from the responses to the odor. In Figure 2B, sensilla were classified as having two neurons if two different spike amplitudes were automatically detected and/or corrected responses to the fruit odor mix were above 20 Hz.

### RNA-sequencing data analysis

Purity-filtered reads were adapters- and quality-trimmed with Cutadapt (version 1.8 (Martin, 2011)). Reads matching to ribosomal RNA sequences were removed with fastq_screen (version 0.11.1). Remaining reads were further filtered for low complexity with reaper (version 15-065) (Davis et al., 2013). Reads were aligned to the *Drosophila melanogaster* BDGP6.92 genome using STAR (version 2.5.3a (Dobin et al., 2013)). The number of read counts per gene locus was summarized with htseq-count (v. 0.9.1) (Anders et al., 2015) using *Drosophila melanogaster*.BDGP6.92 gene annotation. The quality of the RNA-seq data alignment was assessed using RSeQC (v. 2.3.7) (Wang et al., 2012).

Statistical analysis was performed for genes in R (version 3.5.3). Genes with low counts were filtered out according to the rule of one count per million in at least one sample. Library sizes were scaled using TMM normalization (EdgeR package version 3.24.3) (Robinson et al., 2010) and log-transformed with limma cpm function (Limma package version 3.38.3) (Ritchie et al., 2015).

Differential expression was computed with limma for paired samples by fitting the six samples into a linear model and performing the comparison PCD-blocked antennae versus controls. Comparison of read number for *GFP* (encoded by the *Or49a-GFP* transgene) was performed by mapping reads to the *GFP* sequence with Bowtie2 (Langmead and Salzberg, 2012): control antennal RNA: 139±9.5 reads/sample (mean ± standard deviation); PCD-blocked antennal RNA: 227±8.7 reads/sample.

Moderated t-test was used for the comparison on a subset of 83 expressed *D. melanogaster* genes including: *Or, lr* and *Gr* genes as well as the four pro-apoptotic genes (*grim*, *rpr*, *hid* and *skl*). For multiple testing correction, the *p*-values were adjusted by the Benjamini-Hochberg method, which controls the false discovery rate (Benjamini and Hochberg, 1995). The volcano plot (Figure 3A) was generated in R by plotting the log_2_(fold change PCD-blocked vs control) against the -log(p value). Data points were shaded according to mean expression value across all samples.

### Statistics

Statistical analyses and plotting were made in RStudio (v1.1.463 R Foundation for Statistical Computing, Vienna, Austria, 2005; R-project-org), except for the RNA-seq analyses (described above). For statistical analyses, normality was first assessed on datasets using a Shapiro test. If both datasets were normally distributed, a two-sided t-test was performed; otherwise, a Wilcoxon-rank sum test was performed.

## DATA AND CODE AVAILABILITY

Raw data and custom-made analysis scripts are available upon request.

## RESOURCES

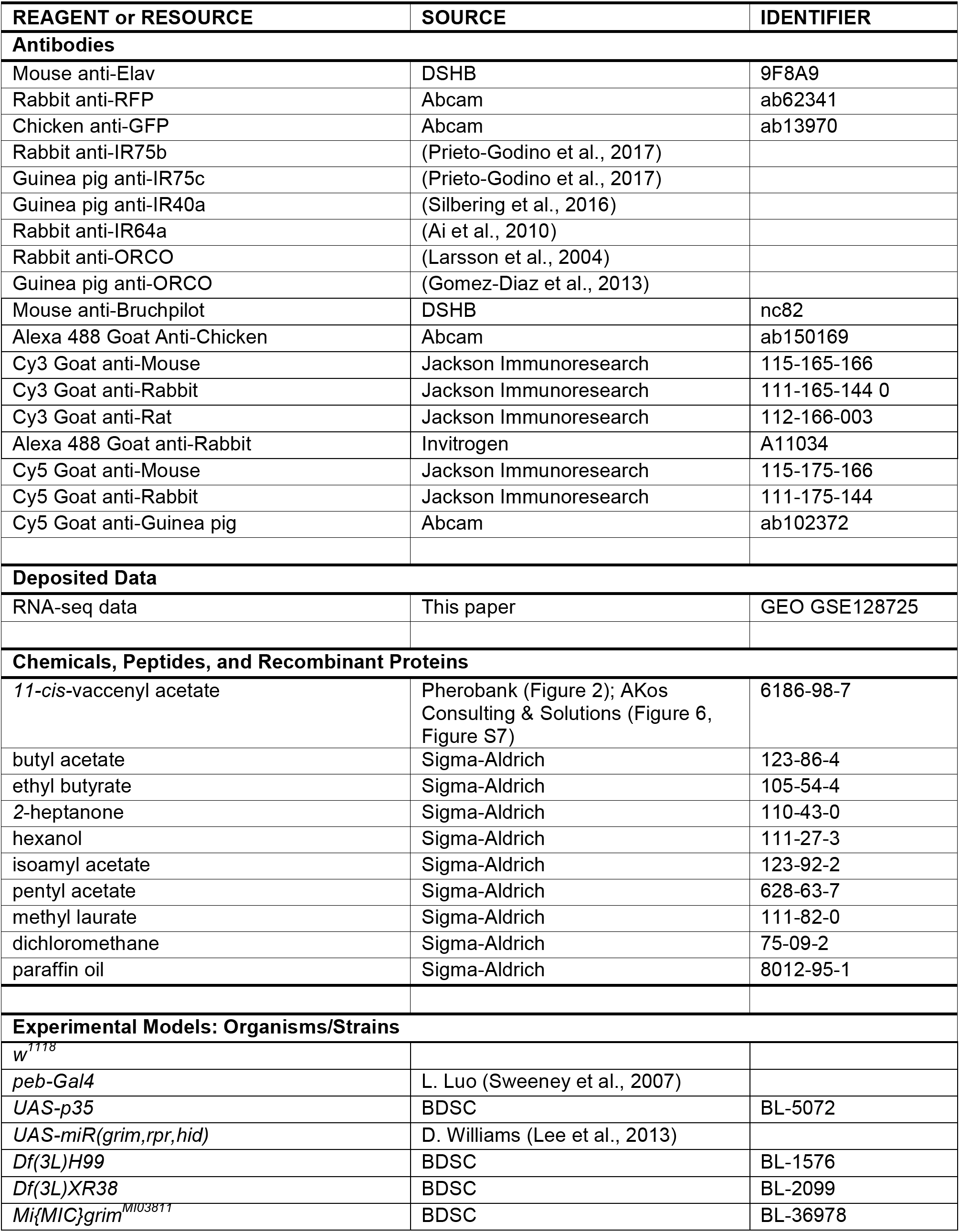
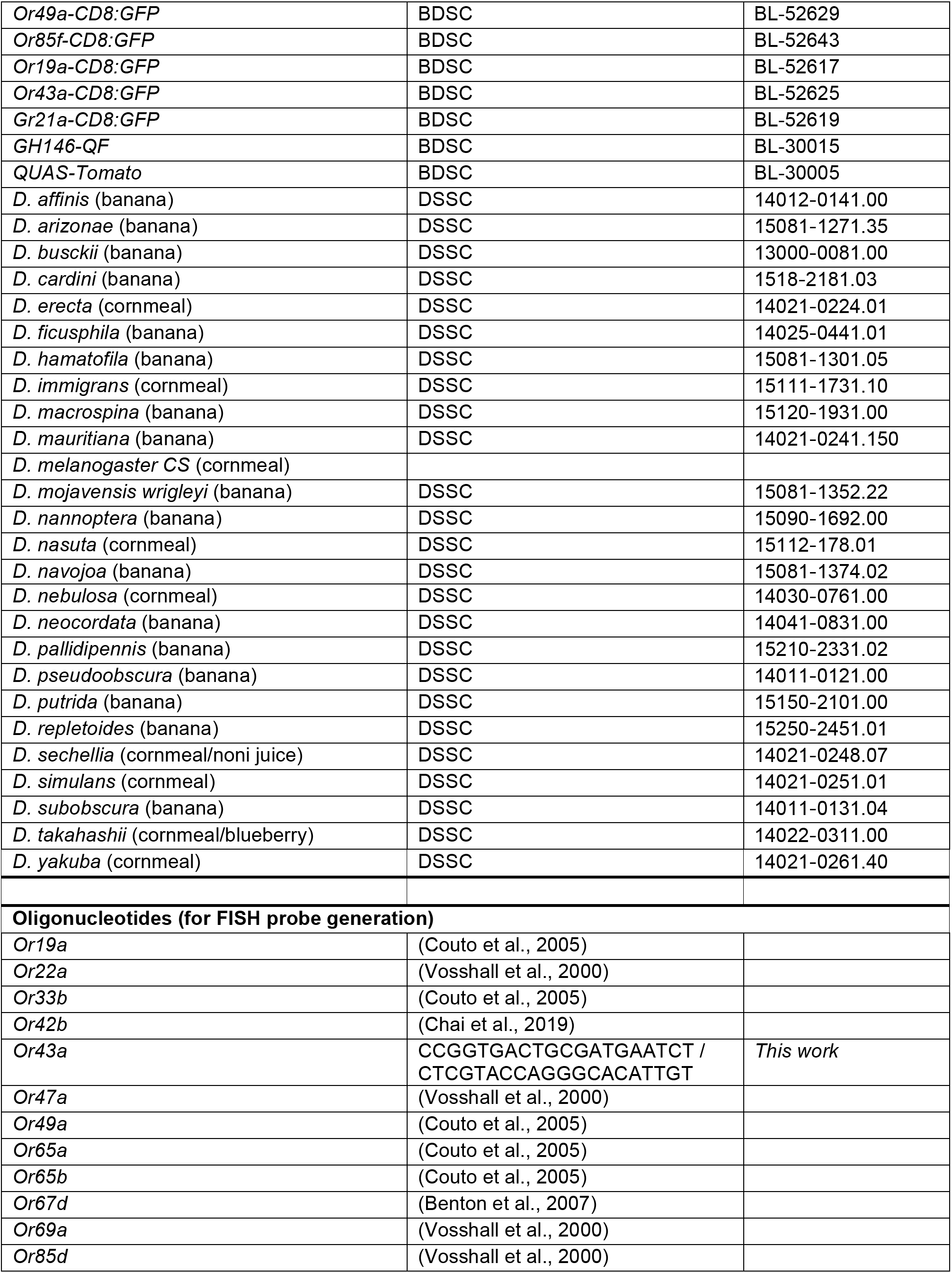
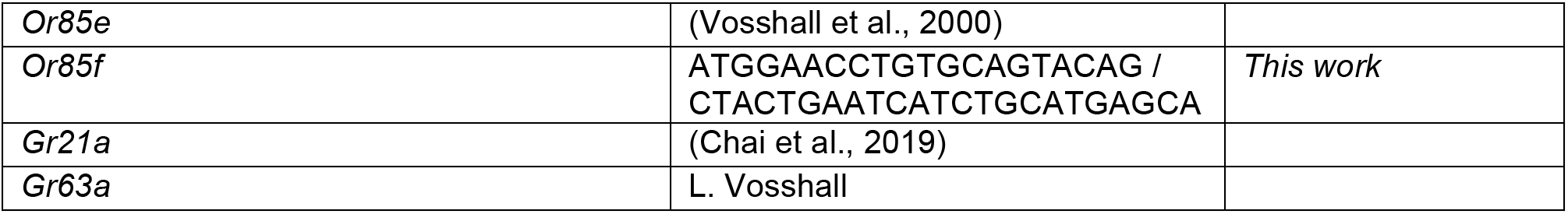

