## Supplementary Information for "Functional integration of “undead” neurons in the olfactory system"

**Figure S1. Automated quantification of Elav-positive olfactory sensory neurons.**

Representative example of Elav expression in whole-mount antennae from control (*peb-Gal4/+*) and PCD-blocked (*peb-Gal4/+;UAS-p35/+*) animals. Middle: output of automated image segmentation of the same antennae used for quantification of OSN number (see Methods). Right: overlay of both images. Scale bar = 10 μm.

**Figure S2. Undead neurons express a subset of olfactory receptor genes.**

**(A)** Representative images of RNA FISH or anti-GFP immunostaining for the indicated *Or* genes in whole mount antennae of control (*peb-Gal4/+*) and PCD-blocked (*peb-Gal4/+;UAS-p35/+*) animals. Scale bar = 10 μm. Quantifications of neuron numbers are shown at the bottom. *** indicates *Or49a*-GFP $p = 1.444 \times 10^{-12}$ (t-test) (n = 14 and 15 (control and PCD-blocked, respectively)), Or65a $p = 4.701 \times 10^{-6}$ (Wilcoxon-sum rank test) (n = 14 and 15), Or69a $p = 1.811 \times 10^{-5}$ (Wilcoxon-sum rank test) (n = 13 and n = 15), Or85f p = 0.01375 (t-test) (n = 16 and n = 10).

**(B)** Representative images of RNA FISH for the indicated *Or* genes in whole mount antennae of control (*peb-Gal4/+*) and PCD-blocked (*peb-Gal4/+;UAS-p35/+*) animals. Scale bar = 10 μm. Quantifications of neuron numbers are shown at the bottom. ns indicate Or35a p = 0.7132 (t-test) (n = 9 and 8), Or67d p = 0.05341 (Wilcoxon-sum rank test) (n = 5 and 13), Gr63*a* p = 0.30325 (Wilcoxon-sum rank test) (n = 10 and n = 10), Ir40a p = 0.763 (t-test) (n = 6 and 10).

**Figure S3. Undead neurons can be found in novel, reproducible locations.**

Additional representative images of RNA FISH for the indicated *Or* genes, anti-GFP immunostaining of the *Or49a*-GFP reporter, or anti-IR64a immunostaining in whole mount antennae of control (*peb-Gal4/+* (or *peb-Gal4/+*;*Or49a-GFP/+*)) and PCD-blocked (*peb-Gal4/+;UAS-p35/+* (or *peb-Gal4/+*;*Or49a-GFP/UAS-p35*)) animals. Scale bar = 10 μm. The pink dashed lines encircle cells in the PCD-blocked antennae that express the corresponding receptors outside their usual spatial domain.

**Figure S4. *Or*-GFP reporters faithfully recapitulate receptor expression.**

**(A)** Representative images of anti-GFP immunofluorescence in whole mount antennae of control (*peb-Gal4/+;;Or19a-GFP/+* and *peb-Gal4/+;Or43a-GFP/+*)) and PCD-blocked (*peb-Gal4/+;UAS-p35/+;Or19a-GFP/+* and *peb-Gal4/+;Or43a-GFP/UAS-p35*) animals. Scale bar = 10 μm. Quantifications of neuron numbers are shown at the bottom. *** indicates *Or19a*-GFP $p = 8.009 \times 10^{-6}$ (t-test) (n = 13 and 15 (control and PCD-blocked, respectively)), *Or43a*-GFP $p = 5.432 \times 10^{-8}$ (t-test) (n = 10 and 10). The pink dashed line encircles cells in PCD-blocked antennae that express the corresponding *Or*s outside their usual spatial domain.

**(B)** Representative images of anti-GFP immunostaining and RNA FISH for *Or85f* in whole mount antennae of control (*peb-Gal4/+;;Or85f-GFP/+*) and PCD-blocked (*peb-Gal4/+;UAS-p35/+;Or85f-GFP/+*) animals. Scale bar = 10 μm. Co-expression quantifications are shown at the bottom. * indicates *Or85f* mRNA p = 0.01375 (t-test) and *Or85f*-GFP p = 0.0153 (t-test) (n = 9 and 9).

**Figure S5. Blocking cell death produces extra glomeruli-like structures in the antennal lobe.**

Representative images of nc82 immunofluorescence in whole mount brains of control (*peb-Gal4/+;;grim*$^{MI03811(EGFP)}$/+) and PCD-blocked (*peb-Gal4/+;UAS-p35/+;grim*$^{MI03811(EGFP)}$/+) animals. Equivalent glomeruli found in control and PCD blocked brains are labeled for reference with dashed blue lines; potential novel glomerular structures (or displaced original glomeruli) in PCD-blocked animals are shown with dashed pink lines. Scale bar = 10 μm.

**Figure S6. Undead olfactory sensory neurons form novel wiring properties in the brain.**

**(A)** Additional representative images of combined anti-GFP and nc82 immunofluorescence in whole mount brains of control (*peb-Gal4/+;Or49a-GFP/Or49a-GFP*, *peb-Gal4/+;;Or19a-GFP/+*, *peb-Gal4/+;Or43a-GFP/+*) and PCD-blocked (*peb-Gal4/+;Or49a-GFP/Or49a-GFP,UAS-p35*, *peb-Gal4/+;UAS-p35/+;Or19a-GFP/+*, *peb-Gal4/+;Or43a-GFP/UAS-p35*) animals. Scale bar = 10 μm.

**(B)** Additional representative images of combined anti-GFP, anti-RFP and nc82 immunofluorescence in whole mount brains of control (*peb-Gal4/+;Or49a-GFP/Or49a-GFP;GH146-QF,QUAS-Tomato/+*, *peb-Gal4/+;;GH146-QF,QUAS-Tomato/Or19a-GFP*, *peb-Gal4/+;Or43a-GFP/+;GH146-QF,QUAS-Tomato/+*) and PCD-blocked (*peb-Gal4/+;Or49a-GFP/Or49a-GFP,UAS-p35;GH146-QF,QUAS-Tomato/+*, *peb-Gal4/+;UAS-p35/+;GH146-QF,QUAS-Tomato/Or19a-GFP*, *peb-Gal4/+;UAS-p35/Or43a-GFP;GH146-QF,QUAS-Tomato/+*) animals. Only the relevant planes are shown. Scale bar = 10 μm.

**Figure S7. Electrophysiological distinction of at1 and at4 sensilla.**

Representative traces of extracellular recordings of neuronal responses to a 0.5 s pulse (black horizontal bar) of methyl laurate (diluted 1:10 v/v), cVA (1:10) or solvent (dichloromethane) in at1 or at4 sensilla of *D. melanogaster* (n = 5) and *D. nasuta* (n = 5). Methyl laurate permits functional distinction of these sensillum classes, as it does not activate the Or67d neuron in *D. melanogaster*, or either neuron in the 2-neuron at1 sensilla of *D. nasuta*. By contrast, this pheromone robustly activates at4 sensilla neurons (corresponding to the Or47b and Or88a OSN classes in *D. melanogaster* (Dweck et al., 2015)).

**Figure S8. Projections of *Gr21a*-GFP-expressing undead neurons from the maxillary palps.**

Additional representative images of combined anti-GFP and nc82 immunofluorescence in whole mount brains of control (*peb-Gal4/+;Gr21a-GFP/+*) and PCD-blocked (*peb-Gal4/+;Gr21a-GFP/UAS-p35*) animals with excised antennae. Both antennal lobes are shown for each brain, showing the contralateral projections of undead palp neurons expressing *Gr21a*-GFP. Scale bar = 10 μm.

Figure S1

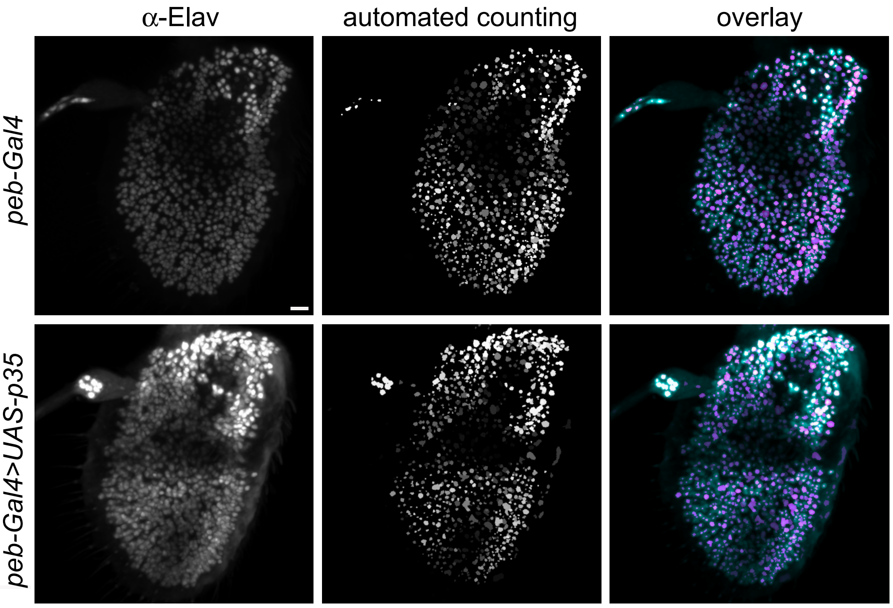

Figure S2

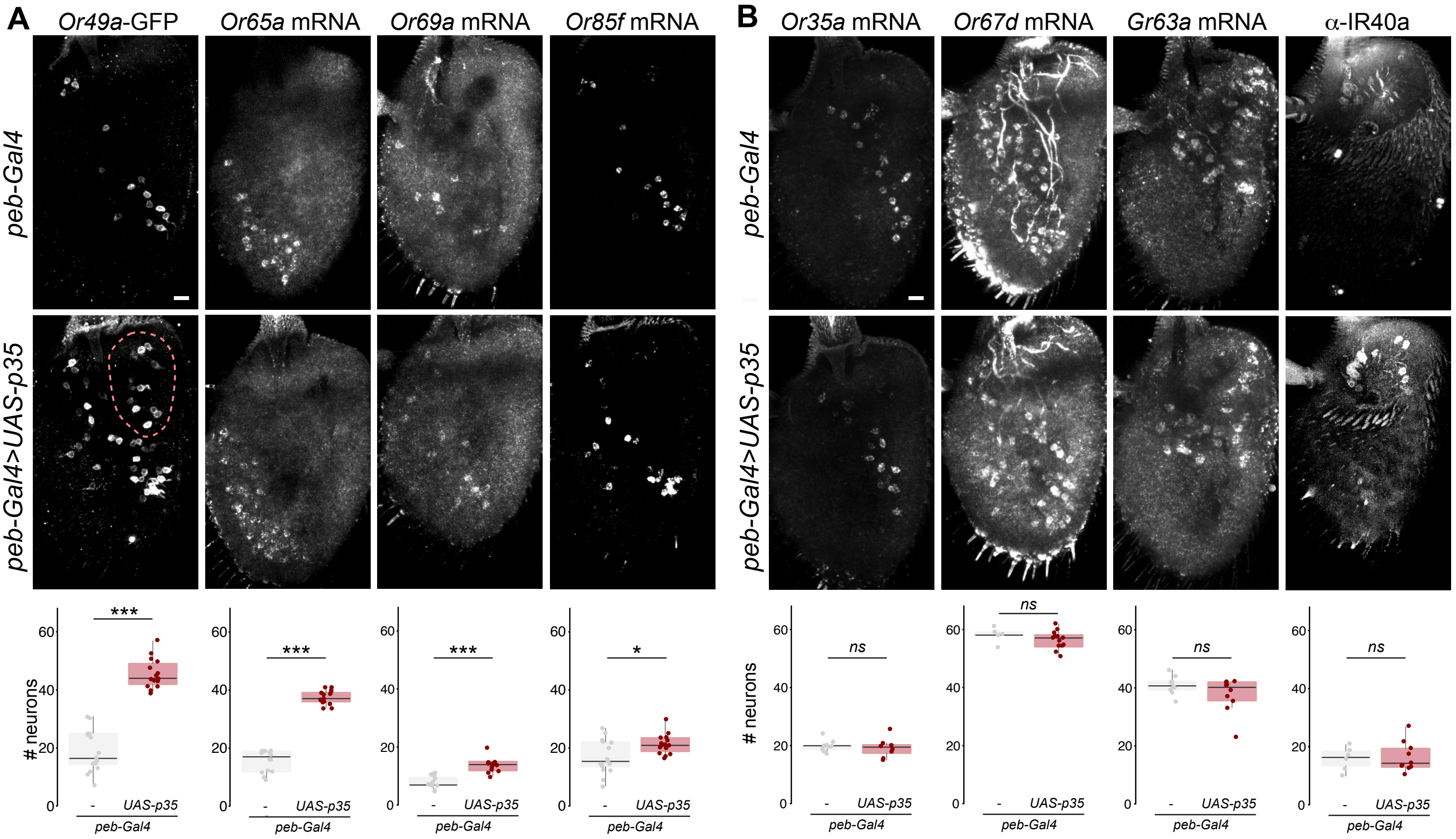
A
Or49a-GFP
Or65a mRNA
Or69a mRNA
Or85f mRNA
peb-Gal4
peb-Gal4>UAS-p35
### neurons
***
***
***
*
-
UAS-p35
peb-Gal4
B
Or35a mRNA
Or67d mRNA
Gr63a mRNA
α-IR40a
ns
0
20
40
60

Figure S3

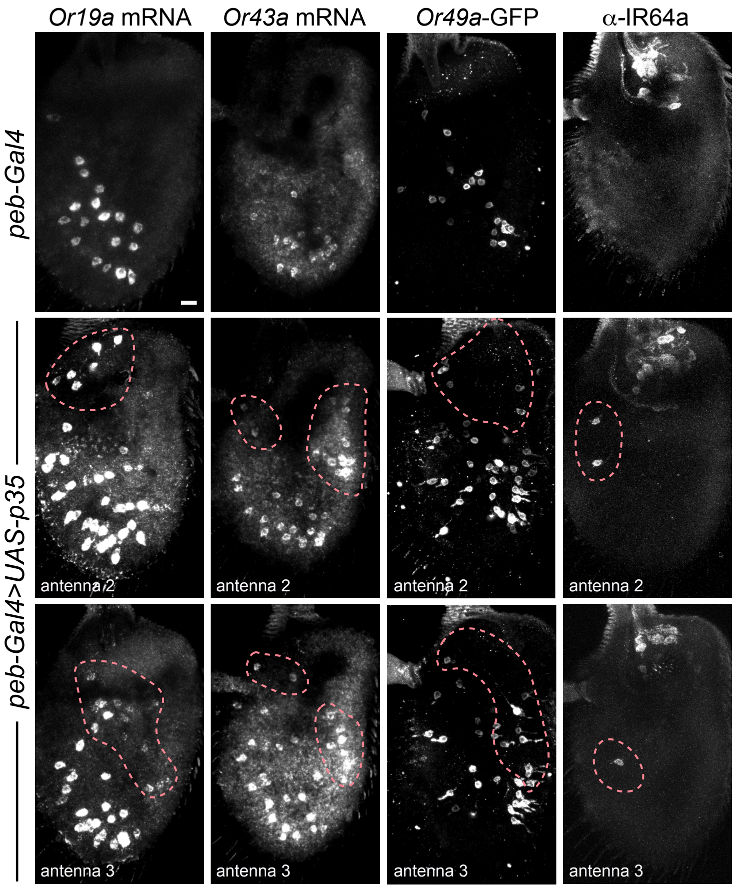

Or19a mRNA
Or43a mRNA
Or49a-GFP
α-IR64a
peb-Gal4
peb-Gal4>UAS-p35
antenna 2
antenna 2
antenna 2
antenna 2
antenna 3
antenna 3
antenna 3
antenna 3

Figure S4

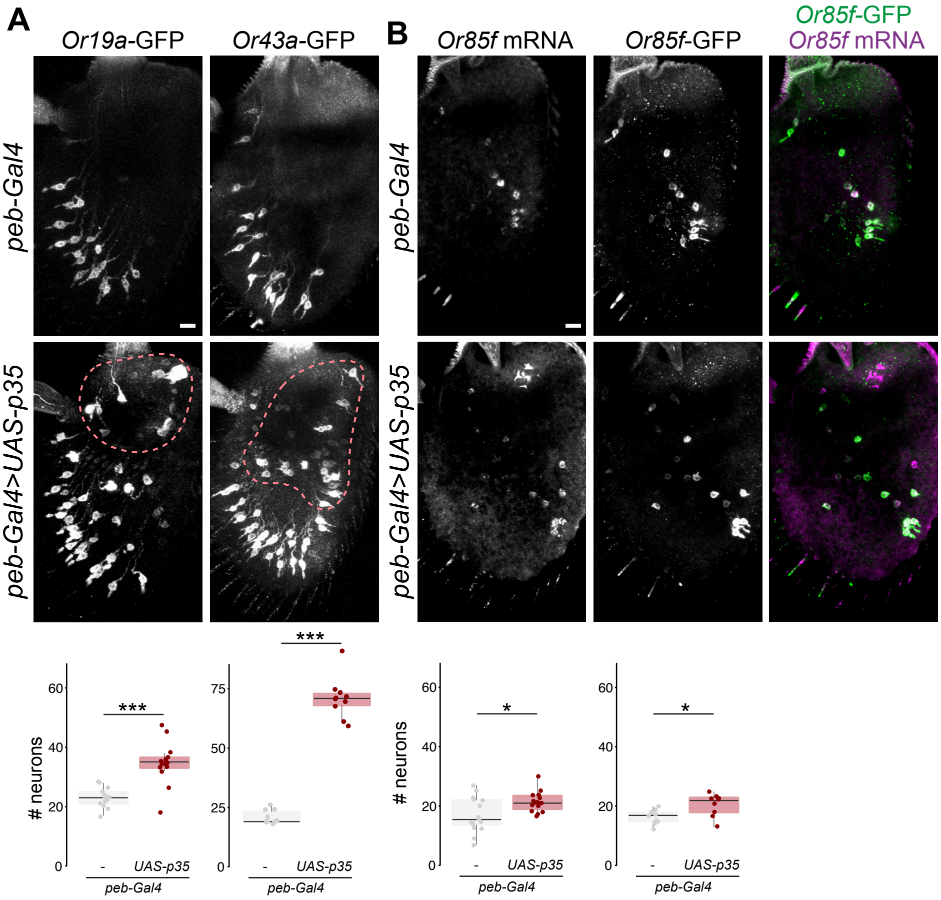

A
Or19a-GFP
Or43a-GFP
B
Or85f mRNA
Or85f-GFP
Or85f-GFP
Or85f mRNA
peb-Gal4
peb-Gal4>UAS-p35
peb-Gal4
peb-Gal4>UAS-p35
### neurons
***
***
*
*
-
UAS-p35
peb-Gal4

Figure S5

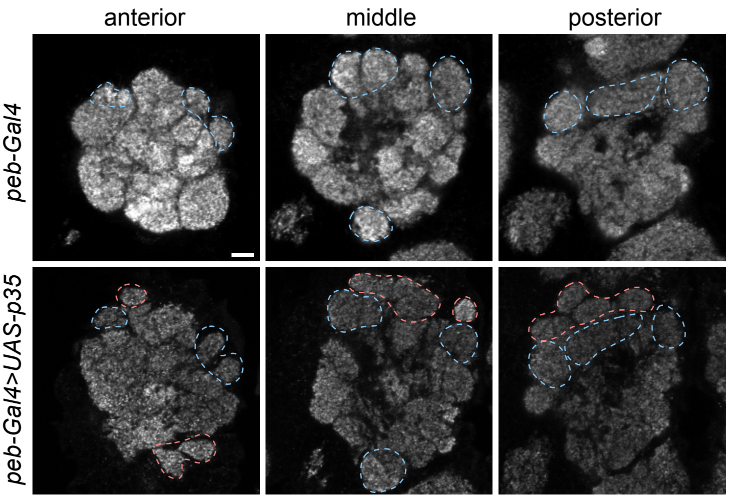

Figure S6

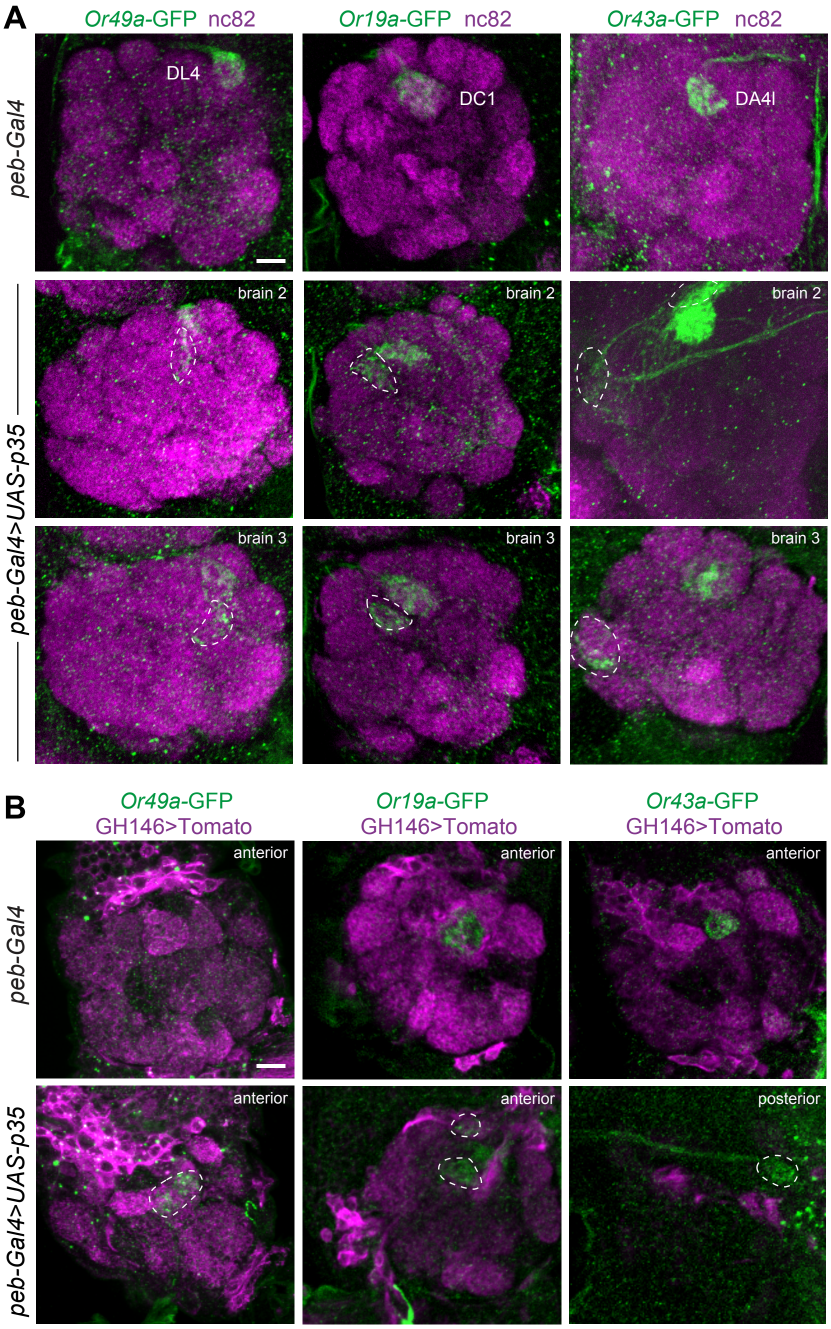

A
Or49a-GFP nc82
Or19a-GFP nc82
Or43a-GFP nc82
peb-Gal4
DL4
DC1
DA4l
peb-Gal4>UAS-p35
brain 2
brain 2
brain 2
brain 3
brain 3
brain 3
B
Or49a-GFP
GH146>Tomato
Or19a-GFP
GH146>Tomato
Or43a-GFP
GH146>Tomato
peb-Gal4
anterior
anterior
anterior
peb-Gal4>UAS-p35
anterior
anterior
posterior

Figure S7

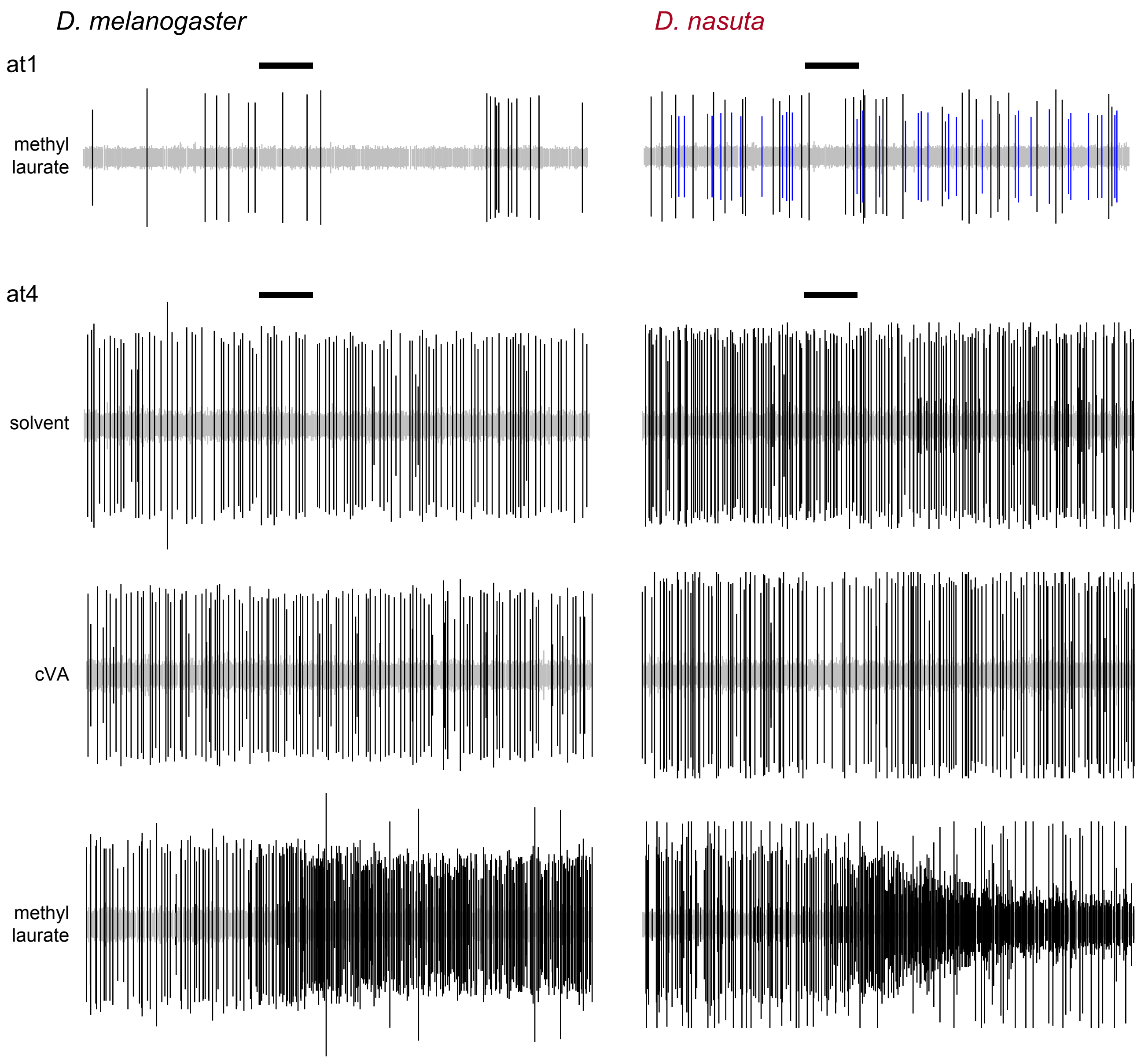
D. melanogaster
D. nasuta
at1
methyl laurate
at4
solvent
cVA
methyl laurate

Figure S8

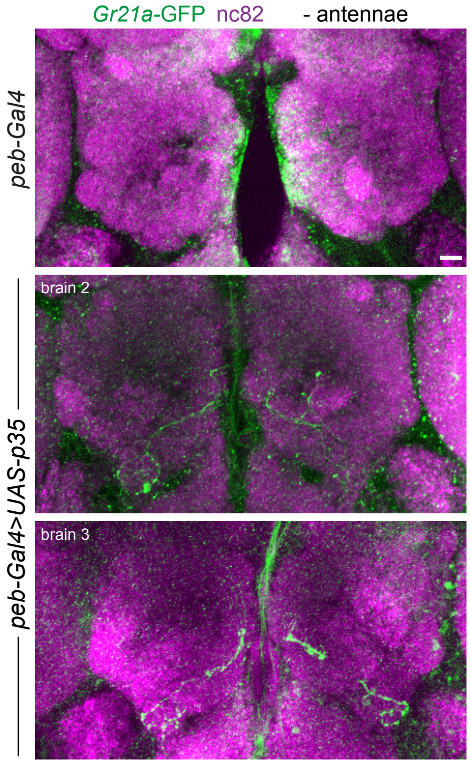

*Gr21a*-GFP nc82
- antennae
*peb-Gal4*
brain 2
brain 3
*peb-Gal4>UAS-p35*
